# Seasonal plasticity of symbiotic strategies clarifies coral holobiont resistance and resilience

**DOI:** 10.1101/2025.11.20.689604

**Authors:** Ariana S. Huffmyer, Emma L. Strand, Serena Hackerott, Kevin H. Wong, Danielle M. Becker, Dennis Conetta, Kristina X. Terpis, Ferdinand Pfab, Juliet M. Wong, Zoe Dellaert, Francis J. Oliaro, Ross Cunning, Jose M. Eirin-Lopez, Steven B. Roberts, Roger M. Nisbet, Hollie M Putnam

## Abstract

As coral reefs face unprecedented declines driven by thermal stress and the breakdown of coral symbiosis (i.e., coral bleaching), restoration efforts increasingly rely on coral health and resilience rankings. However, seasonal plasticity in symbiosis and holobiont metabolism, along with the presence of cryptic species, can complicate data interpretation. Therefore, quantifying seasonal plasticity in coral physiology and incorporating genetic identification are essential for accurately interpreting and drawing conclusions from trait-based and fitness-based analyses. To test the effect of seasonal and site variation on physiological plasticity we sampled three ecologically dominant genera, *Acropora*, *Pocillopora*, and *Porites* across three lagoon sites (n=15 tagged colonies genus^−1^ site^−1^) on the north shore of Mo’orea French Polynesia in January, March, September, and December of 2020. We identified coral host and intracellular symbiotic Symbiodiniaceae to the highest taxonomic resolution possible and quantified 13 physiological variables within the holobiont. Genetic analyses identified *A. pulchra* along with cryptic lineages in *Pocillopora* (*P. meandrina, P. tuahiniensis*) and *Porites* (*P. evermanni, P. lobata/lutea*). *A. pulchra* was dominated by *Durusdinium trenchii* and also contained *Symbiodinium microadriaticum.* Symbiont communities differed between cryptic congeners, with *P. meandrina* hosting *Cladocopium latusorum* and *P. tuahiniensis* hosting *Cladocopium pacificum*, whereas *P. evermanni* and *P. lobata/lutea* both hosted *Cladocopium* (C15), but each with unique C15 profiles. Weedy taxa such as *Acropora* and *Pocillopora* displayed a cycle of symbiont boom and bust in response to seasonally variable light and temperature, likely contributing to the high stress sensitivity of these taxa. In contrast, despite seasonal environmental variability, *Porites* displayed greater symbiont stability, with temperature—rather than light—serving as the stronger explanatory variable of seasonal variation in host physiology. Increased host biomass under cooler conditions, which provides energy reserves, may serve as an important stabilizing factor in massive *Porites* well-documented stress resilience. Collectively, our data provide essential evidence of the need for integrative analyses considering baseline physiological states across seasons along with host and symbiont genetics, particularly in light of the plethora of climate change related stress test assays taking place throughout the year across coral taxa with cryptic lineages.

## Introduction

Coral reefs are among the most ecologically diverse ecosystems in the world and at their foundation are coral holobionts: meta-organisms composed of the coral host, algal symbiont, and associated microbiome (Goulet et al., 2020). These partners exchange crucial nutrients to sustain a mutually beneficial symbiotic relationship (Ainsworth et al., 2010; Roth, 2014). However, with ocean waters warming rapidly (Oliver et al., 2021), the maintenance of this symbiosis is threatened (van Woesik et al., 2022), as temperature-driven metabolic changes destabilize the partnership and increase the risk of dysbiosis, bleaching, and mortality. With increasingly frequent mass coral mortality events (Hughes et al., 2017, 2018), there is an immediate need to understand the mechanisms of coral stress response and acclimatization to environmental change (Putnam, 2021). Such shifting baselines in coral physiology (Wall et al., 2021) will impact the range of acclimatization potential possible under future climate change (Grottoli et al., 2014; Logan et al., 2014).

Critical aspects for understanding acclimatization dynamics and organismal stress thresholds include the characterization of the baseline/current physiological state (Cunning & Baker, 2013), the breadth of seasonal variation in trait plasticity (Thornhill et al., 2011), and the genetic identity of the holobiont partners (Berkelmans & van Oppen, 2006). In particular, there is growing evidence that the multifaceted combination of environmental conditions at any point in the year has the potential to modulate a coral’s bleaching susceptibility and response to stress (Holcomb et al., 2012; Scheufen, Krämer, et al., 2017). Such examples demonstrate that both coral and endosymbiont metabolism and physiology are responsive to seasonal environmental change. For example, variation across season is seen in symbiont density (Fitt et al., 2000), tissue biomass (Gómez et al., 2023; Thornhill et al., 2011; Trumbauer et al., 2021; van de Water et al., 2018), skeletal growth (Edmunds & Putnam, 2020; Gómez et al., 2023; van de Water et al., 2018), gametogenesis (Keith et al., 2016), lipid content and stable carbon and nitrogen isotopes (Trumbauer et al., 2021), photophysiology (Scheufen, Iglesias-Prieto, et al., 2017; Trumbauer et al., 2021; Ulstrup et al., 2008; van de Water et al., 2018; Warner et al., 2002), microbial community associations (Miller & Bentlage, 2024; Sharp et al., 2017; van de Water et al., 2018), gene expression (Wuitchik et al., 2019), and the epigenome (Hackerott et al., 2023; Rodríguez-Casariego et al., 2020). Thus, it is critical to investigate the magnitude and mechanisms of these effects across common and dominant reef building taxa under natural seasonal variations, especially as novel climate change conditions exacerbate seasonal heatwaves.

Coral function is a product of the tight metabolic interactions between the coral host, their algal intracellular symbionts, and diverse associated microbial communities (i.e., bacteria, fungi, archaea) (Boilard et al., 2020; M. Li et al., 2023; Radecker et al., 2015; Siboni et al., 2008; Thompson et al., 2014). As holobionts, reef-building corals benefit from symbiotic associations with microbial communities, but also contend with the vulnerabilities of their partners. In particular, the nutritional relationship between the coral host and intracellular symbionts (family Symbiodiniaceae; hereafter referred to as “symbionts” or “Symbiodiniaceae”) requires a balance in nutritional cycling to maintain stable relationships, especially under stress (Rädecker et al., 2021, 2023). There is clear evidence that the identity of the Symbiodiniaceae and their photophysiological characteristics (Wall et al., 2020) have functional implications for the holobiont in terms of carbon translocation (Stat et al., 2008), skeletal growth (Little et al., 2004), depth acclimatization (Ziegler et al., 2015), and the sensitivity of the bleaching response (Cunning et al., 2016; Hoadley et al., 2019; Roach et al., 2021). In a notable example, the performance of pairs of the same species of coral (*Montipora capitata*) living immediately adjacent to each other is distinctly different when the colonies are dominated by symbionts in the genus *Cladocopium* or *Durusdinium*. Between these holobionts, there are distinct phenotypes of bleaching response (Cunning et al., 2016), carbon translocation from symbiont to host (Allen-Waller & Barott, 2023), metabolomic profiles (Roach et al., 2021), growth (Walker et al., 2025) and bleaching-induced partial mortality (Matsuda et al., 2020). Therefore, it is critical to evaluate coral holobiont response to environmental variation, considering the identity of both the host and the symbiotic partners (Bellantuono et al., 2012).

Coral reefs in Mo’orea, French Polynesia, the location of our study, experience dynamic environmental conditions, with seasonal and diel temperature changes of ∼3-4°C ((Edmunds et al., 2010), current study), variable nutrient levels (∼1.6 fold seasonal change in nitrogen *Turbinaria* tissue content) (Adam et al., 2021), and spatio-temporal variation in physical oceanographic dynamics (Edmunds et al., 2010; Leichter et al., 2012). While many coral taxa thrive in these dynamic environments, the genera *Porites*, *Pocillopora*, and *Acropora* have remained dominant in the back reefs (Dahl & Edmunds, 2024), each with distinct life history characteristics, symbiotic associations (Putnam et al., 2012), and cryptic species (Burgess et al., 2021; Forsman et al., 2015; Johnston et al., 2018; Rassmussen et al., 2025). *Porites* are mounding corals with thick tissues (Barnes & Lough, 1992; Edmunds et al., 2012; Yost et al., 2013) that pass symbionts through vertical transmission (i.e., symbionts are passed to offspring through the oocyte) and are considered environmentally tolerant (Loya et al., 2001; Putnam et al., 2012). *Pocillopora* are branching corals with thin tissues (Yost et al., 2013) that pass symbionts through vertical transmission and exhibit high recruitment rates after a disturbance (Grigg & Maragos, 1974; Holbrook et al., 2018) despite their high sensitivity to environmental disturbance (Burgess et al., 2021; Loya et al., 2001). *Acropora* are also branching corals with relatively thin tissues (Bucher & Harrison, 2018) that pass symbionts via horizontal transmission and exhibit high sensitivity to environmental change and stress (Loya et al., 2001). Further, *Pocillopora* and *Acropora* spp. (complex clade (Romano & Cairns, 2000)) are symbiotic generalists hosting multiple species/strains of *Cladocopium* spp., *Symbiodinium* spp. and/or *Durusdinium* spp. (Putnam et al., 2012). In contrast, *Porites* spp. (robust clade (Romano & Cairns, 2000)) are symbiotic specialists, primarily hosting one species or a set of closely related *Cladocopium* spp. (e.g., C15) (Putnam et al., 2012). Variability in species’ symbiotic associations, environmental sensitivity, and tissue and skeletal morphology provide a framework in which to test the dynamics of physiological plasticity across spatial and temporal gradients in important reef-building corals. Further, understanding cryptic lineage variation within genera is essential because physiological and ecological differences among genetically distinct, but morphologically similar, lineages can influence interpretations of species resilience (Burgess et al., 2021; Forsman et al., 2015; Johnston et al., 2018) and the mechanisms that underpin coral persistence in changing environments.

In light of their different life history strategies and symbiotic associations, we chose these three dominant genera of reef builders (*Porites*, *Pocillopora*, and *Acropora)* to assess holobiont physiological responses across spatiotemporal environmental variation. We conducted extensive physiological sampling of corals from three environmentally variable lagoon sites in Mo’orea, French Polynesia, across four time points in one year (January, March, September, and November 2020). Given the presence of cryptic species in *Acropora* spp. (Rassmussen et al., 2025), *Pocillopora* spp. (Burgess et al., 2021; Johnston et al., 2018), and *Porites* spp. (Forsman et al., 2015), we conducted genetic sequencing and determined that samples collected included *A. pulchra* as well as representatives of *Pocillopora tuahiniensis, Pocillopora meandrina, Porites evermanni,* and *Porites lobata/lutea*. Therefore, we examined the physiological responses to site and seasonal variation, considering cryptic host lineages and their associated symbiont communities.

## Materials and Methods

### A. Replication statement

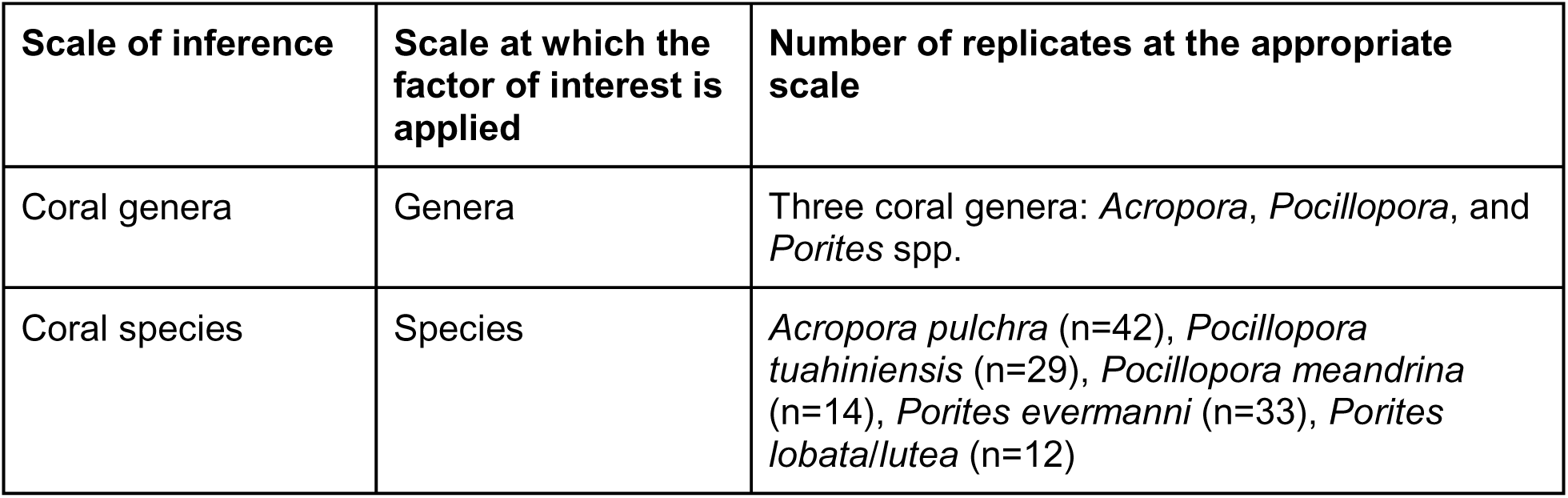
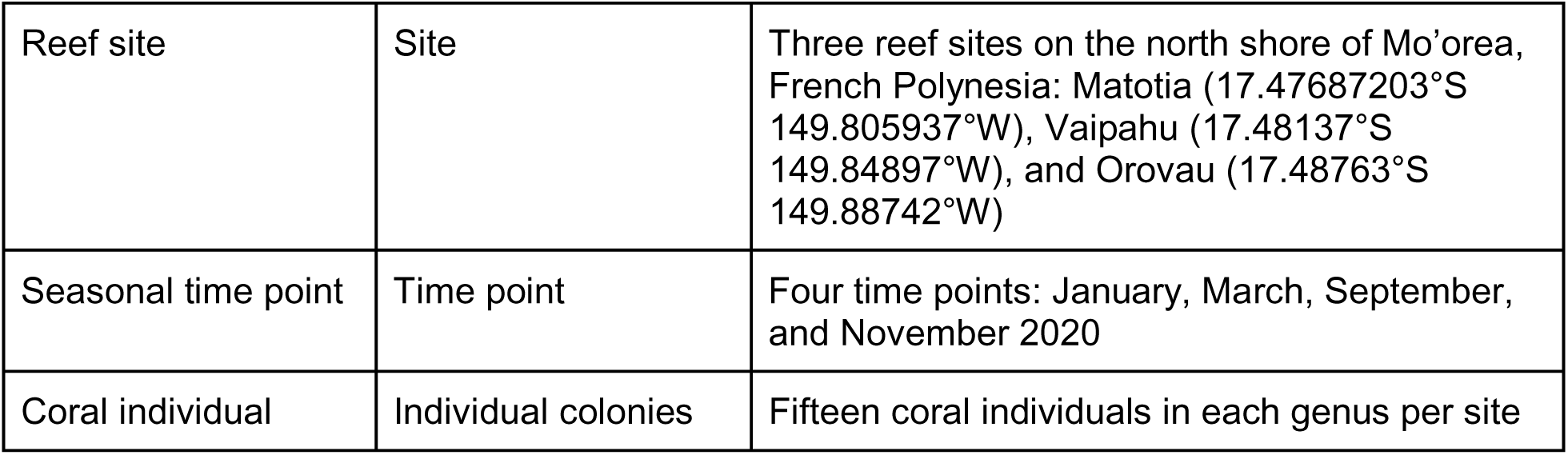

| Scale of inference | Scale at which the factor of interest is applied | Number of replicates at the appropriate scale |
| --- | --- | --- |
| Coral genera | Genera | Three coral genera: *Acropora*, *Pocillopora*, and *Porites* spp. |
| Coral species | Species | *Acropora pulchra* (n=42), *Pocillopora tuahiniensis* (n=29), *Pocillopora meandrina* (n=14), *Porites evermanni* (n=33), *Porites lobata/lutea* (n=12) |
| Reef site | Site | Three reef sites on the north shore of Mo'orea, French Polynesia: Matotia (17.47687203°S 149.805937°W), Vaipahu (17.48137°S 149.84897°W), and Orovau (17.48763°S 149.88742°W) |
| Seasonal time point | Time point | Four time points: January, March, September, and November 2020 |
| Coral individual | Individual colonies | Fifteen coral individuals in each genus per site |

### B. Site and genus selection

We tracked physiological traits of three coral genera (*Porites* spp., *Pocillopora* spp., *Acropora* spp.) across spatial and temporal gradients on the north shore backreef of Mo’orea, French Polynesia. Fifteen *Acropora* coral genets were fragmented from the lagoon on the north shore (17.48393°S, 149.83405°W), individually tagged, and transplanted to each site (n=15 per site) in October 2019. Individual genets of *Pocillopora* and *Porites* were identified and tagged at the three sites in October 2019 (n=15 per genus per site).

Three study sites were selected across an environmental gradient based on previously characterized nutrient conditions (nitrogen tissue content in *Turbinaria* macroalgae) (Adam et al., 2021). The sites selected were Orovau (relatively lower N:C; 17.48763°S 149.88742°W), Vaipahu (relatively middle N:C; 17.48137°S 149.84897°W), and Matotia (relatively higher N:C; 17.47687203°S 149.805937°W) (**Fig 1**; site names correspond to the Toponymes de Polynésie Française map from Direction du système d’information de la Polynésie Française). Temperature, pH, and light were measured throughout the study to characterize differences in environmental conditions between these sites (described below). Tagged colonies were sampled at each of the three sites at four time points: January (28 Dec 2019 - 14 Jan 2020), March (27 Feb - 14 March), September (8 Sept - 28 Sept), and November (30 Oct - 21 Nov) 2020. These sampling time points allowed for n=2 samplings during the wet, humid period (January and March 2020) and n=2 samplings during the cooler, drier season (September and November 2020; (Adam et al., 2021)). Of the 135 total colonies tagged and sampled in January 2020, 112 were sampled in March, 99 in September, and 107 in November. Reduced colony sample size at subsequent time points was due to colony mortality or inability to locate colonies. All experimental and field operations took place at the University of California, Berkeley, Richard B. Gump South Pacific Research Station (UCB Gump Station).

**Fig 1.**
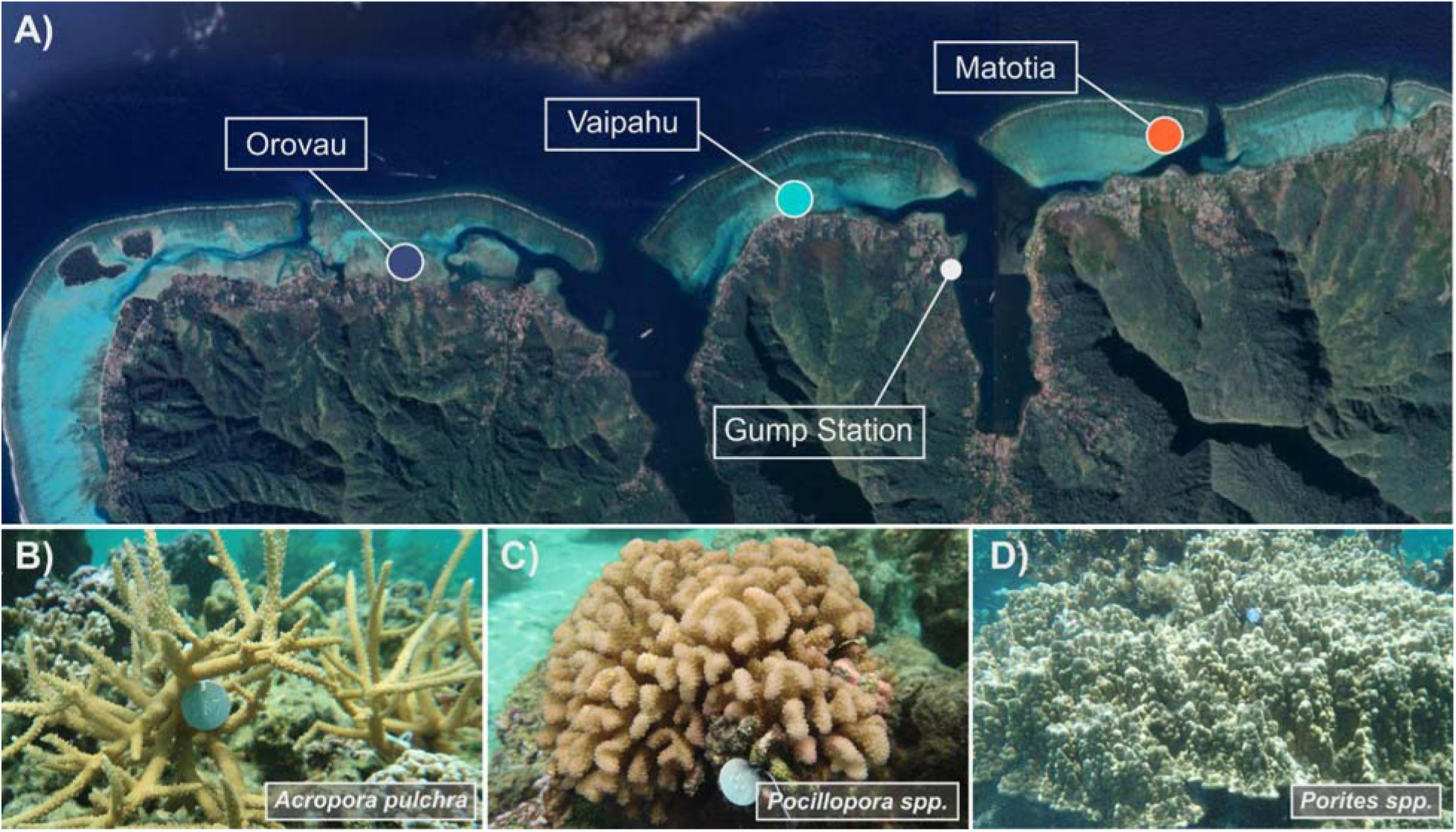
Study design. (A) Three sites (Orovau, Vaipahu, Matotia) were established in October 2019 by transplanting in 10 colonies of (B) *Acropora pulchra* and tagging 15 colonies of C) *Pocillopora* spp. and D) *Porites* spp. at each site. Colonies were sampled in January, March, September, and November of 2020.

### C. Environmental characterization

Environmental data loggers were deployed on the reef at each study site in October 2019 and data were collected during each sampling time point. HOBO v2 U22 Temperature loggers (U22-001; resolution = 0.02°C; accuracy = ± 0.21°C) were calibrated with a Digital Traceable Thermometer (4000CC; accuracy = ± 0.05°C; resolution = 0.0001°C) prior to deployment. HOBO MX pH loggers (MX2501; accuracy and resolution = ± 0.20 mV) were calibrated with pH 7.00 and pH 4.01 NBS standards prior to deployment at each time point. HOBO U24 Conductivity Loggers (U24-002-C; resolution = 2 µS/cm) were calibrated with Conductivity Solution RICCA Cat #2248 Conductivity Standard 50,000 µS/cm at 25°C. Odyssey Xtreem PAR loggers were calibrated to a LICOR cosine Underwater Quantum Sensor (LI-192).

Due to errors in logging and damaged or missing loggers (when COVID in 2020 reduced our travel capacity to the field), light, pH, and salinity were only available for brief time windows, and therefore we focus primarily on temperature data from the study sites, which is available from the entire time series. Temperature data was collected from November 2019 through November 2020 (**Fig 2A**). To provide additional environmental context for each sampling time point, we examined publicly available data through the Mo’orea Coral Reef Long Term Ecological Research site (Washburn & Brooks, 2024). We visualized data for solar radiation (kWh m^−2^) and cumulative rainfall (mm). These data were collected from one station on the north shore of the island of Mo’orea and therefore we cannot distinguish differences in these characteristics between study sites. For each time point, we calculated the mean (± st. dev.) temperature (**Fig 2B**), light (**Fig 2C**) and rainfall (**Fig 2D**) as the mean of daily values for all observations in the 4 weeks prior to sampling for each time point. Data was not available for the months of February or March 2020 for light or rainfall and therefore we calculated means for the month of April to approximate conditions during the March sampling time point.

**Fig 2.**
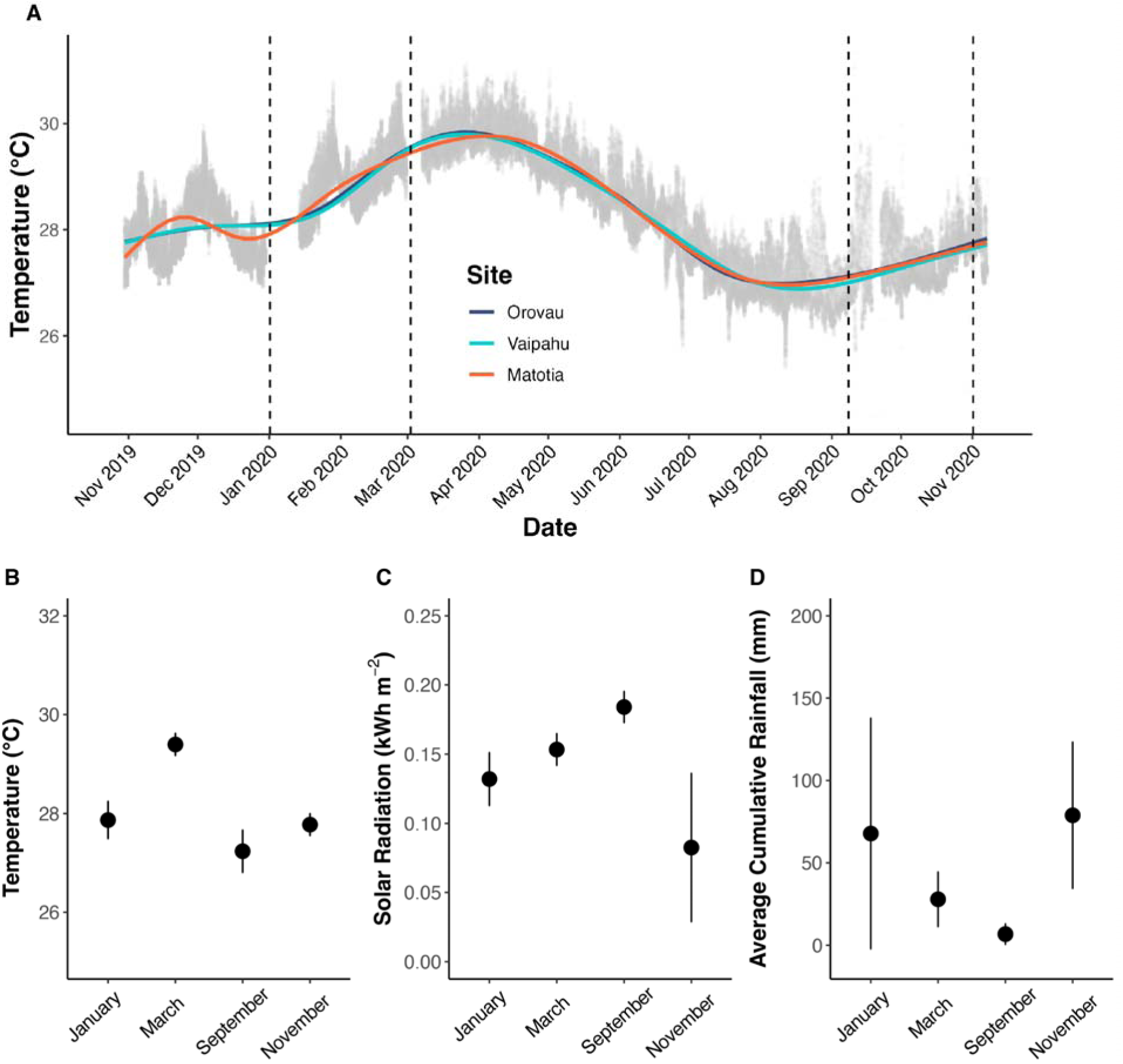
Environmental variables. (A) Temperature (°C) at each site across the one-year time series study. Generalized additive model lines shown for each site (Orovau = blue; Vaipahu = cyan; Matotia = orange). Temperature observations from data loggers are shown in gray. Dashed lines indicate sampling time points (Jan, Mar, Sep, and Nov 2020). (B) Water temperature (°C) as measured in the present study averaged for the 4 weeks prior to sampling time point. (C) Solar radiation (kWh m^−2^) averaged for the 4 weeks prior to sampling time point. (D) Rainfall accumulation (mm) averaged for the 4 weeks prior to sampling time point. Solar radiation and rainfall data obtained from MCR LTER (Washburn & Brooks, 2024). All values show mean ± st. dev.

### D. Sampling and live sample processing

At each time point, tagged colonies were sampled by clipping three fragments. Two 2 cm biopsies were preserved for molecular analyses by immediately snap-freezing in liquid nitrogen on a boat. Samples were returned to University of California Berkeley Richard B. Gump South Pacific Research Station where they were clipped into 1.5 mL tubes with 600 µL of DNA/RNA Shield (Zymo #R1100-250) followed by storage at −40°C until they were transported to URI for processing (CITES FR2198700194-E). One 3 cm fragment was collected live from each colony returned to the UCB Gump station and placed in flow-through seawater tables for 1-6 days during physiological sampling with an equal number of samples per day per genus randomly selected from the water tables for processing. Sample sizes are presented in **Table S1**. Here, we provide a general description of each metric and refer readers to **Supporting Information** for further details.

#### 1. Photosynthesis-irradiance (PI) curves

Photosynthesis-irradiance (PI) curves were characterized using oxygen production and consumption rates with oxygen measurements of coral fragments under increasing light levels (10 min each at 0 [dark], 18, 68, 113, 169, 243, 499, 709, 844, and 1025 µmol photons m^2^ s^1^ photosynthetically active radiation; PAR). Oxygen evolution rates were calculated for each light level interval using localized linear regressions (alpha=0.2, percentile rank) in the *LoLinR* package (Olito et al., 2017), normalized to surface area, and calculated as µmol O_2_ cm^−2^ h^−1^. PI curves were generated using the quadratic equation (Falkowski & Raven, 2013) to calculate maximal photosynthetic rate (P_MllX_), apparent quantum yield (AQY), dark respiration (R_D_), saturating irradiance (J_K_) and compensation irradiance (J_C_) for each colony at each time point. See **Supporting Information** for details.

#### 2. Instantaneous calcification rates

Instantaneous calcification rate for each fragment was measured using the total alkalinity (TA) anomaly technique (Chisholm & Gattuso, 1991). Fragments were incubated for 90 min at 28°C and 500 µmol photons m^−2^ s^−1^ PAR (above saturating irradiance; see **Supporting Information**) with water samples collected from the initial water source and following the incubation (*n*=8 coral and *n*=2 blank samples) and preserved with 75 μL of 50% saturated mercuric chloride (HgCl_2_) solution. Total alkalinity was measured by following the open cell potentiometric titration (SOP3b; Dickson et al., 2007). See **Supporting Information** for details. Calcification rates were calculated following the total alkalinity (TA) anomaly technique (Chisholm & Gattuso, 1991). TA values were normalized to salinity, blank corrected, and normalized to surface area to calculate calcification rates as µmol CaCO_3_ cm^−2^ hr^−1^. Following incubation, samples were flash frozen in liquid nitrogen for physiological analyses.

#### 3. Physiology sample processing

Coral tissue was separated from the skeleton using an Iwata Eclipse HP-BCS airbrush with cold 1X Phosphate Buffer Saline (PBS) solution, resulting in a tissue slurry. Tissue slurry was homogenized using a sterilized PRO Scientific Bio-Gen PRO200 Homogenizer. Two 1 mL aliquots of tissue homogenate were centrifuged at 13,000 rpm for 3 min (Eppendorf Centrifuge 5415D). The resulting symbiont cell pellet from one aliquot was stored at −40°C for symbiont biomass analyses. The host coral supernatant of the second aliquot was removed and stored at −40°C for host biomass, protein, and antioxidant analyses. The remaining algal pellet in the second aliquot was resuspended in 1 mL of ice-cold 1X PBS. 500 µL of the resuspended pellet was stored in a −40°C freezer until processing for symbiont density counts and another 500 µL was stored at −40°C for chlorophyll concentration determination. The airbrushed coral skeletons were placed in a drying oven (Fisher Scientific Isotemp Oven) at 60°C for 4 h.

Surface area (cm^2^) was measured on dried skeletons using the wax-dipping method (Stimson & Kinzie, 1991; Veal et al., 2010) and biomass was measured on both host and symbiont fractions as ash-free dry weight and normalized to fragment surface area (AFDW; mg AFDW cm^−2^). Host soluble protein was quantified using a Bovine Serum Albumin assay (Thermo Scientific Pierce BCA Protein Assay) and normalized to tissue biomass as mg protein mg AFDW^−1^. Antioxidant capacity (TAC) of the host tissue was measured using the Cell BioLabs OxiSelect TAC Assay Kit (Cat # STA-360) according to the manufacturer’s instructions. TAC was normalized to biomass and calculated as µmol Copper Reducing Equivalents (CRE) per mg AFDW.

For chlorophyll quantification, a 500 µL aliquot was centrifuged at 13,000 rpm for 3 min (Eppendorf Centrifuge 5415D), the supernatant removed, and 1 mL of 100% acetone added for 24 h. Symbiont chlorophyll concentration (*a* and *c_2_* pigments) were quantified via spectroscopy with blank subtraction of 750 nm, and using absorbance at 663 nm and 630 nm, following >equation 3 for dinoflagellates in 100% acetone from (Jeffrey & Humphrey, 1975), with correction for path length. Chl *a* and *c_2_* pigments were normalized to tissue biomass to generate µg pigment mg AFDW^−1^. See **Supporting Information** for details. The sum of *a* and *c_2_* pigments was calculated to generate total chlorophyll pigment (µg pigment mg AFDW^−1^). Chlorophyll content was also normalized to symbiont cell density to generate µg pigment cell^−1^. Symbiodiniaceae cell density (n=6 replicate counts per sample) of resuspended symbiont pellets was measured using an Improved Neubauer Haemocytometer (Marienfeld Superior, Lauda-Königshofen, Germany) on a dissecting microscope and was calculated as cells mg AFDW^−1^.

### E. Molecular characterization

We conducted genetic identification of host haplotypes in *Pocillopora* spp. and *Porites spp. Pocillopora* species were identified by amplifying the mitochondrial open reading frame (mtORF) region as described by (Burgess et al., 2021; Johnston et al., 2018) using primers from (Flot et al., 2008) followed by PocHistone 3 region to distinguish species for mtORF haplotype 1a (*P. meandrina* and *P. eydouxi*) as described in (Johnston et al., 2018). *Porites* species were identified using the coral nuclear histone region spanning H2A to H4 (i.e., H2) (Tisthammer et al., 2020). *Acropora* samples were identified by amplifying the mitochondrial control region (CRf, CO3r) using primers from (Vollmer & Palumbi, 2002). Sequences were aligned and analyzed using Geneious Alignment in GENEIOUS PRIME 2020.2.4. See **Supporting Information** for details.

Symbiodiniaceae communities were characterized by metabarcoding of the ITS2 rDNA region (Davies et al., 2022). Symbiodiniaceae ITS2 was amplified using the SYM_VAR primers (Hume et al., 2018) with barcodes and Illumina adapters added following (Kozich et al., 2013). Pooled libraries were sequenced on an Illumina MiSeq using a 500-cycle v2 reagent kit (250 bp paired-end reads) and custom sequencing primers to initiate forward, reverse and index reads (Kozich et al., 2013). See **Supporting Information** for details.

### F. Statistical analysis

#### 1. Univariate analysis

All downstream data analysis was conducted using R Statistical Programming v4.2.2 (R Core Team, 2022). Univariate statistical analyses for each response (log-transformed) were conducted using linear mixed effect models (LMM) within each genus in the *lme4* package (Bates et al., 2015). Time, site, and their interaction were included as main effects with colony as a random intercept to account for repeated measures. For *Pocillopora* spp. and *Porites* spp. that had multiple host haplotypes detected (i.e., “holobionts”), we used colony nested within holobiont nested within genus as a random intercept. Main effects were evaluated using Type III ANOVA tests in the *lmerTest* package (Kuznetsova et al., 2015) with post hoc comparisons using the *emmeans* package (Lenth, 2018). The assumption of residual normality was assessed using quantile-quantile plots. Significance of random effects was tested using ANOVA-like tables in the *lmerTest* package (Kuznetsova et al., 2015).

#### 2. Multivariate analysis

We used a principal component analysis (PCA) to visualize multivariate physiology using all responses (log-transformed) for each genus and then for each holobiont within each genus. Differences in multivariate physiology between groups (i.e., genus or holobiont) were tested using a permutation multivariate analysis of variance (PERMANOVA) and dispersion (PERMDISP) using euclidean distance in the *vegan* package (Okansen & et al., 2017). Analyses were conducted at the level of combined holobiont responses (all host and symbiont responses) as well as host and symbiont responses (**Fig 3**). We then conducted multivariate analyses within each genus, including site, time point, and their interaction as main effects and conducted post hoc comparisons using pairwise permanova tests (*pairwiseAdonis* package; (Martinez Arbizu, 2017)) and Tukey Honest Significant Difference (HSD) tests (*stats* package; (R Core Team, 2022)), respectively. Effect sizes were calculated as Omega-squared values using the MicEco package (*adonis_OmegaSq* function; Russel, 2021). Within each analysis, the proportion of variance in multivariate physiology explained by each main effect variable (site, time point, and holobiont [cryptic host lineage and associated symbiont community]) individually (i.e., controlling for the influence of the other variable) was quantified with variance partitioning and redundancy analyses (RDA) in the *vegan* package (Okansen & et al., 2017). In cases where site and holobiont were confounded (i.e., only one holobiont was found at one of the sites for *Porites* in our study), the model partitioned site and time point effects into combined effects as individual main effects could not be fully separated. To visualize shifts in multivariate physiology across site and time points, we determined multivariate trajectories by calculating the centroid for each time point x site combination and represented the paths between centroids across time as arrows starting at January 2020 and ending at November 2020.

**Fig 3.**
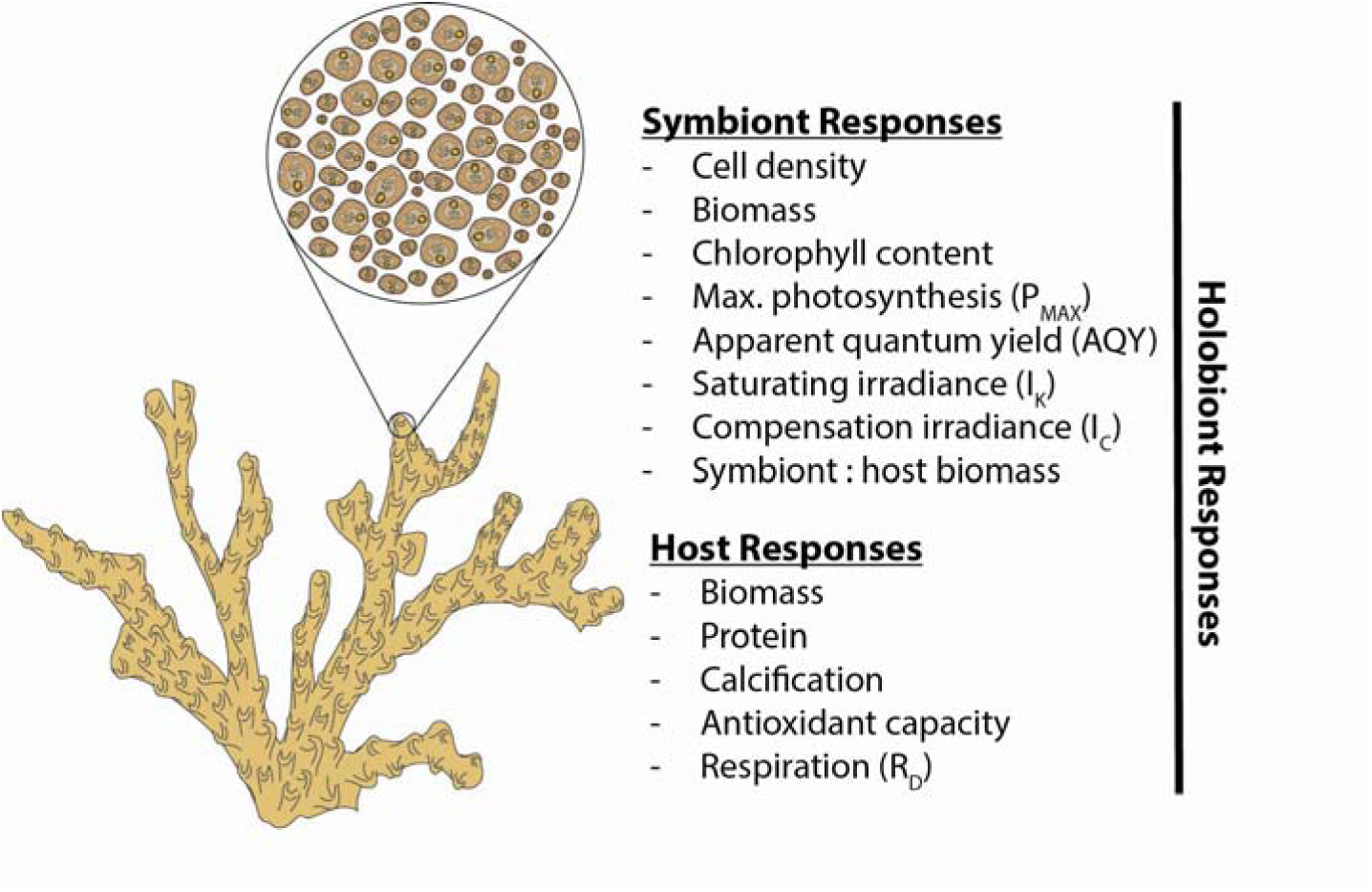
Physiological responses. Physiological response measured in the host and symbiont in this study.

#### 3. Symbiont community ITS2 analysis

Demultiplexed FASTQ files for each sample had the SYM_VAR forward primer prepended to read 1 sequences, and the SYM_VAR reverse primer prepended to read 2 sequences (with dummy quality scores), prior to processing through the SymPortal analysis pipeline (Hume et al., 2019). ITS2 sequences from raw FASTQ files were uploaded and analyzed on the SymPortal remote server (symportal.org; (Hume et al., 2019)). Absolute abundance matrices were analyzed with the *phyloseq* package (v1.44.0; (McMurdie & Holmes, 2013)) to calculate relative abundance and visualize ITS2 Type Profile communities (filtered to remove variants with <1% relative abundance). PERMANOVA statistical tests were performed to assess significant effects of genus, time point, and site on relative abundance of ITS2 profiles using the *vegan* package (Okansen & et al., 2017). Due to significant differences in symbiont communities between genera, we further analyzed the effect of holobiont, site, and time point within each genus (holobiont effects only in *Pocillopora* and *Porites*). Due to the significant effect of genus, two-way ANOVA models were run with site, profile, and holobiont as fixed factors within each genus (holobiont not included for *Acropora*) and Tukey post hoc tests in the *emmeans* package (Lenth, 2018).

We further examined relationships between physiology and variation in the symbiont community through distance-based redundancy analyses (dbRDA), with symbiont community dissimilarity (Bray-Curtis) as the response variable. For each genus, two separate dbRDAs were conducted including the suite of physiological response metrics (**Fig 3**) of either the host or the symbiont as predictor variables. Physiological metrics significantly correlated with variation in the symbiont community were identified with ANOVA-like permutation tests in the *vegan* package (Okansen & et al., 2017).

#### 4. Redundancy analysis of effects of environmental characteristics on physiology

Finally, to examine specific seasonal environmental effects on host and symbiont physiology, we conducted redundancy analyses in the *vegan* package (Okansen & et al., 2017) to model the effects of light and temperature (described above) on host and symbiont physiological responses. Pearson correlation analyses between the environmental conditions identified that rainfall was correlated (P<0.05) with light and temperature and was therefore removed from further analyses. Maximum and minimum light and temperature were also correlated (P<0.05) with the mean value of each metric, and therefore only mean light and temperature were retained for RDA analyses. Mean light and temperature values for each time point were then modeled against the physiological responses of the host and symbiont separately for *A. pulchra, Pocillopora* spp., and *Porites* spp. and visualized as variance explained and in ordination plots.

## Results

### A. Acropora, Pocillopora, and Porites corals exhibit distinct physiological characteristics and cryptic holobiont identities

*Acropora* only contained one host species, *Acropora pulchra* (sensu (Conn et al. 2025); **Fig S1**)*. Pocillopora* and *Porites*, on the other hand, each included two host species. *Pocillopora* colonies were identified as either *Pocillopora meandrina* (*n*=15 total with *n*=9 at Orovau, *n*=3 at Vaipahu, and *n*=3 at Matotia) or *Pocillopora tuahiniensis* (*n*=29 total with *n*=5 at Orovau, *n*=12 at Vaipahu, and *n*=12 at Matotia; **Table S1**; **Fig S1**). *Porites* colonies were identified as either *Porites evermanni* (*n*=33 total with *n*=6 at Orovau, *n*=12 at Vaipahu, and *n*=15 at Matotia) or in the *Porites lobata/lutea clade* (*n*=12 total with *n*=9 at Orovau, *n*=3 at Vaipahu, and *n*=0 at Matotia; **Table S1**; **Fig S1**).

Each genus showed distinct physiological characteristics in the host (PERMANOVA P=0.001; **Fig 4A**) and symbiont (PERMANOVA P=0.001; **Fig 4C**; **Table S2**). Across genera, differences in host physiology were driven by higher antioxidant capacity, calcification, respiration, and host biomass in *Porites* spp. as well as higher AQY, P_MAX_, and symbiont biomass in *Porites* spp. *Acropora pulchra* exhibited greater symbiont cell density, S:H biomass, and chlorophyll (**Fig 4BD**). Multivariate physiology in the host and symbiont were not different between *Pocillopora tuahiniensis* and *Pocillopora meandrina* (**Fig 4AC**; **Table S3**) but symbiont physiology did vary between *Porites evermanni* and *Porites lobata/lutea* (PERMANOVA symbiont P=0.010; host P=0.590; **Fig 4AC**; **Table S3**).

**Fig 4.**
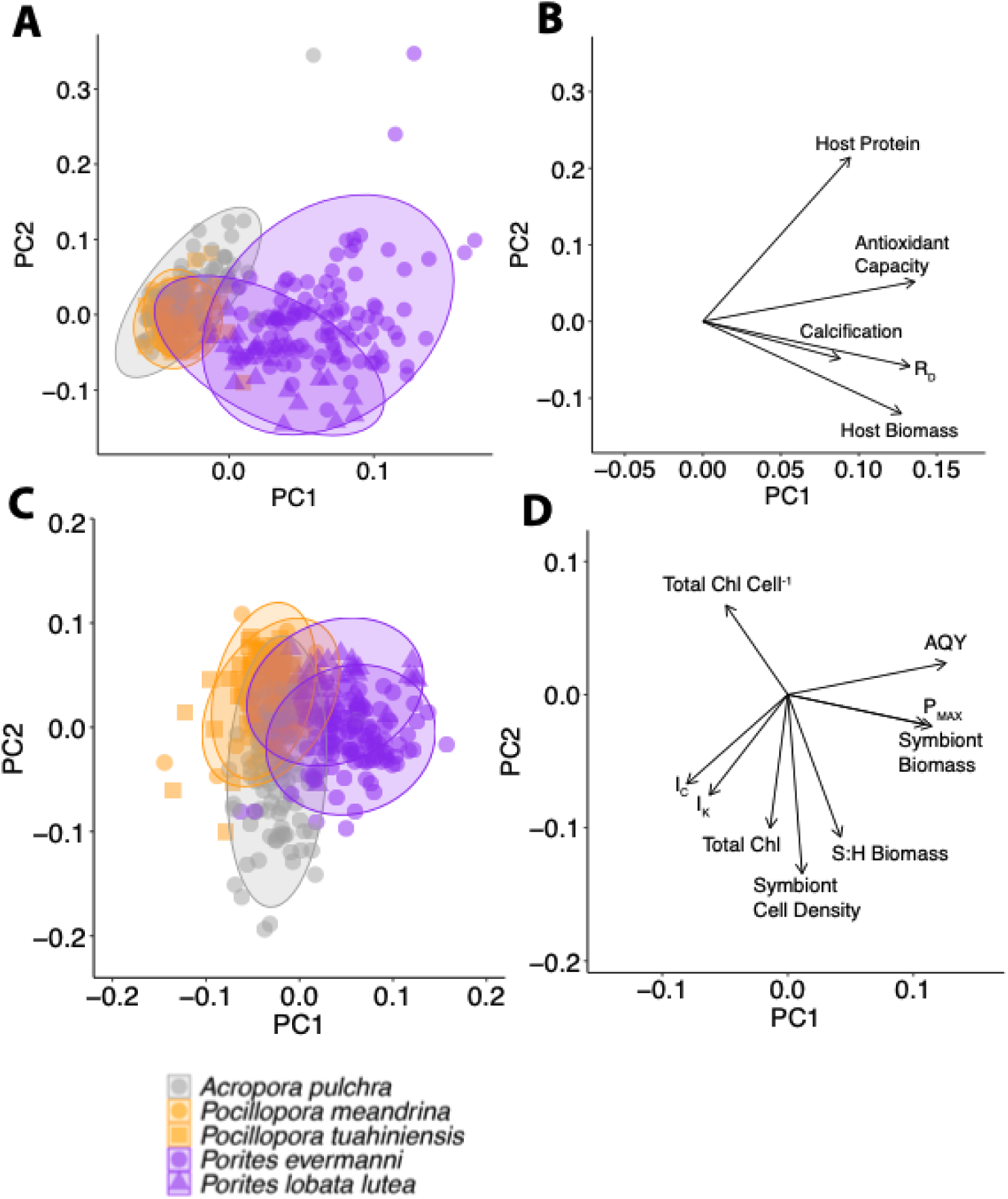
Multivariate physiological responses. (A) Principal components analysis (PCA) of host physiological responses for each holobiont. (B) Biplot of physiological loadings driving separation in multivariate host physiology. (C) PCA of symbiont physiological responses. (D) Biplot of physiological loadings driving separation in multivariate symbiont physiology. Refer to Fig 3 for abbreviations of physiological metrics. In all plots, *Acropora pulchra* is indicated in gray, *Pocillopora* spp. are in orange (*P. meandrina* = circles; *P. tuahiniensis* = squares), and *Porites* spp. are in purple (*P. evermanni* = circles; *P. lobata lutea* = triangles).

Symbiodiniaceae ITS2 profiles did not change over time and were not variable between sites within each genus (**Table S4**). *Acropora pulchra* was associated with a majority (81%) of D1-D1u-D1jb ITS2 profile with lesser relative abundance of three A1 profiles (A1/A1ee, A1/A1ee-A1ep-A1eq, and A1/A1gb-A1ee) (**Fig 5AB**). There were significant differences in symbiont communities between host haplotypes in *Pocillopora* spp. and *Porites spp.* (PERMANOVA P=0.001; **Fig S2**) and differences between haplotypes (PERMANOVA P[profile:holobiont]<0.001; **Table S5**). *Pocillopora meandrina* was associated with *Cladocopium* with 50-60% relative abundance of C42g/C1/C42.2/C42a-C42h-C1b-C42b-C1ew-C42br and lesser component of C1/C42.2/C42g/C42a-C1b-C1au-C1az-C3-C42h and C1d/C1/C42.2/C3-C1b-C3cg-C45c-C115k-C1au-C41p (**Fig 5CD**). In *P. tuahiniensis,* profile C42g/C1/C42.2/C42a-C42h-C1b-C42b-C1ew-C42br had lower relative abundance than in *P. meandrina* and higher presence of C1d-C42.2-C1-C1k-C1b-C3cg and C1d/C42.2/C1/C3cg-C1b-C3cw-C115k-C45c profiles (**Fig 5CD**).

**Fig 5.**
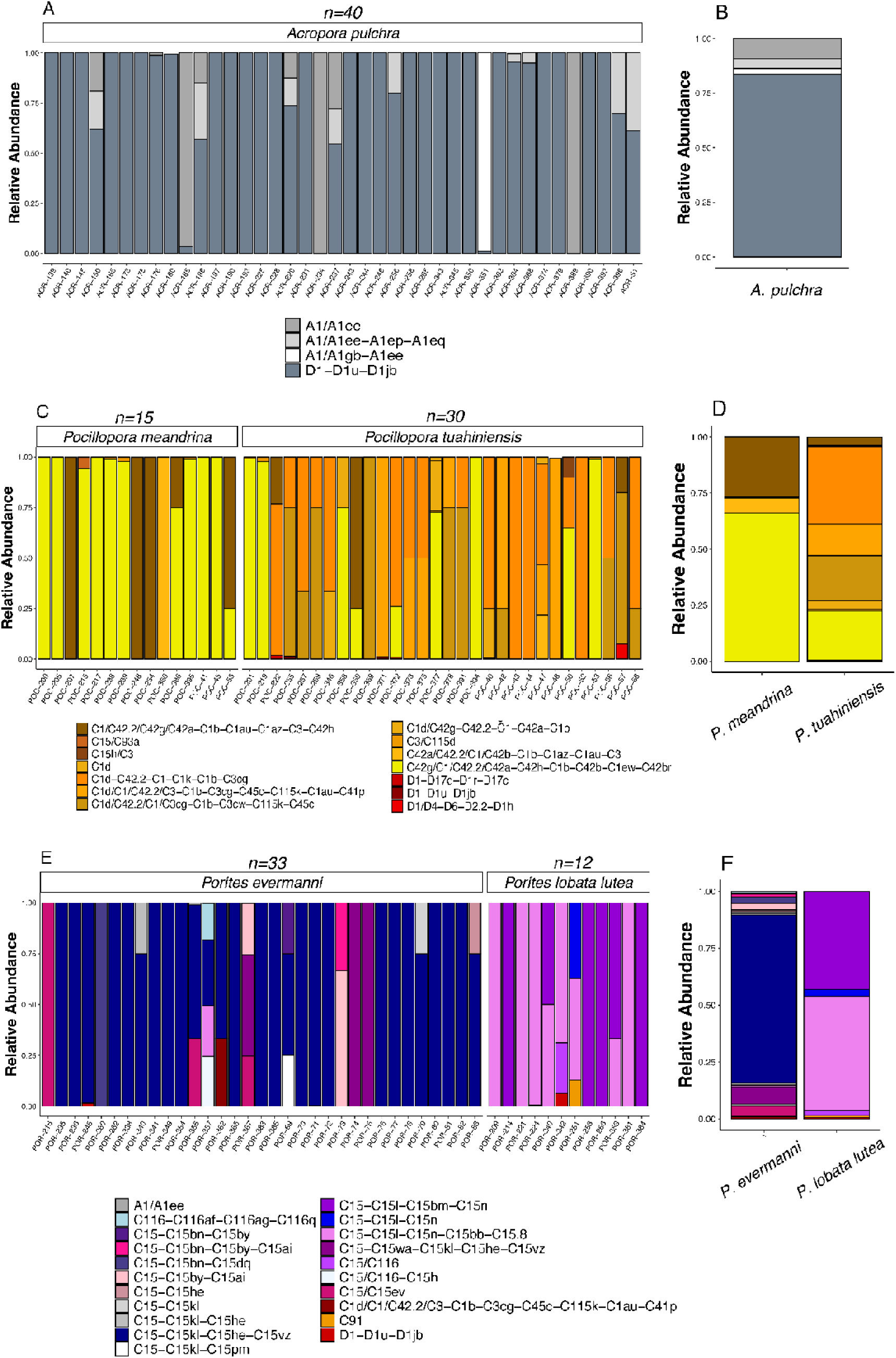
Symbiont community composition. Symbiont ITS2 profile relative abundance for each coral colony in (A) *Acropora pulchra*, (C) *Pocillopora meandrina* and *Pocillopora tuahiniensis*, and (E) *Porites evermanni* and *Porites lobata/lutea*. Mean symbiont ITS2 relative abundance summarized for each holobiont in (B) *Acropora,* (D) *Pocillopora*, and (F) *Porites*. All plots show relative abundance of taxa representing >1% relative abundance of ITS2 profiles calculated at the colony level. ITS2 profile indicated as the majority profile followed by minor profiles in order of decreasing relative abundance separated by dashes and are shown by color. *n* indicates sample size.

*Porites* spp. exhibited clear differences in ITS2 profile relative abundance between host haplotypes. *Porites* spp. showed high fidelity association with *Cladocopium* C15 symbionts but with variation in specific C15 profiles between haplotypes. *Porites evermanni* was associated with a majority of C15-C15kl-C15he-C15vz while *P. lobata/lutea* was associated with C15-C15l-C15n-C15bb-C15.8 and C15-C15l-C15bm-C15n as major components (**Fig 5EF**). Given that each *Pocillopora* and *Porites* haplotype was associated with distinct symbiont communities, we hereafter refer to host and symbiont identity together as distinct “holobionts”. Because there was one host species in *Acropora* (*Acropora pulchra*), there is only one holobiont for *Acropora*.

### B. Strong seasonal effects on physiology

We found significant effects of time point, site, and their interaction on multivariate physiology in both host and symbiont responses (further described below). In all three genera and at each biological level, time point explained the highest proportion of variance of any effect (**Fig 6**, **Table S6**) with significant site, time point and/or interactive effects present in each univariate response (**Table S7**, **Table S8**).

**Fig 6.**
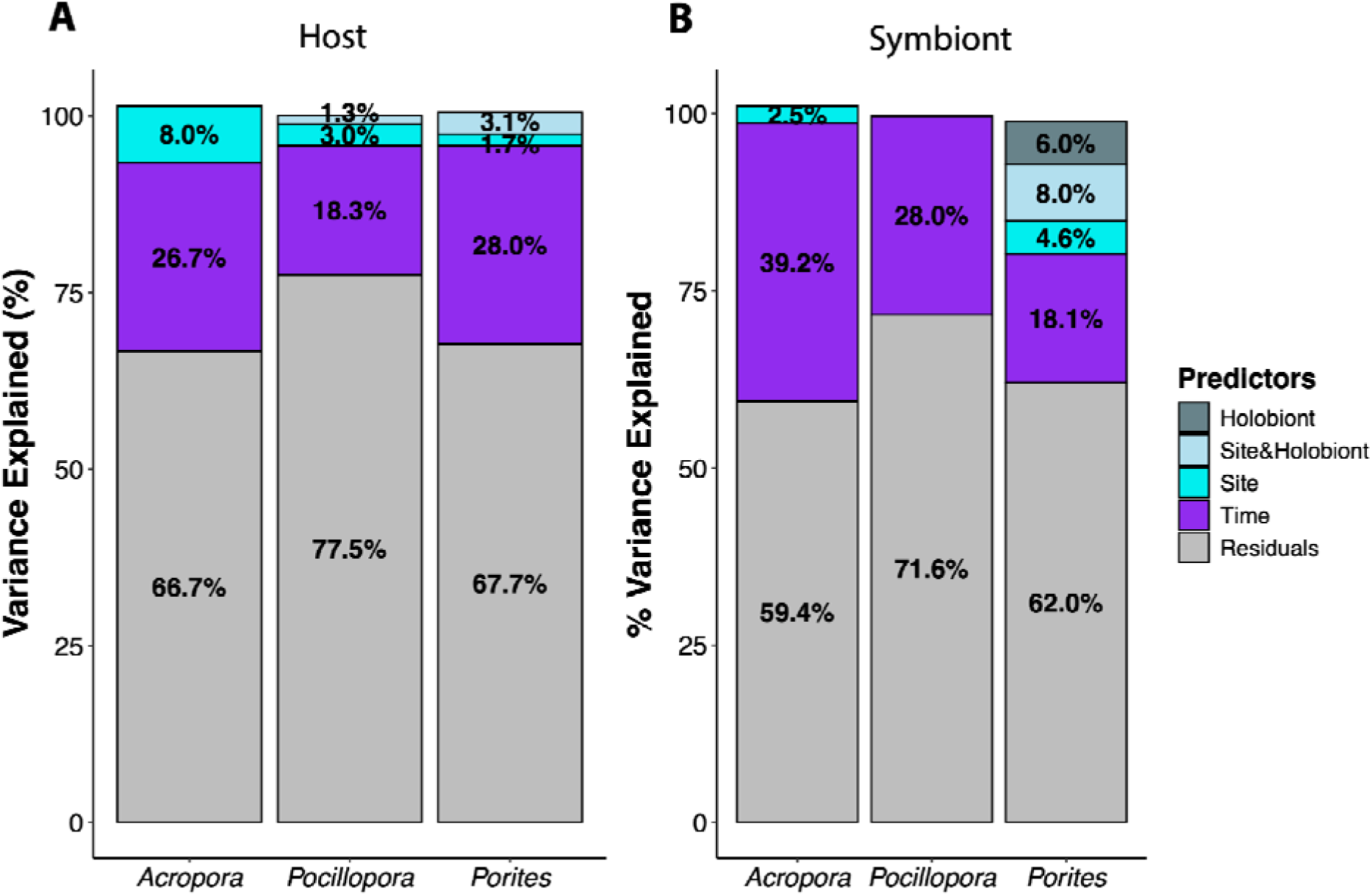
Variance partitioning analysis of effects on physiology. Percent variance explained in multivariate physiology by holobiont (i.e., host species and associated symbiont community; dark gray), combined effects of site and holobiont (light blue), site (cyan) and time point (purple) using redundancy analyses (RDA). Effects shown for host (A), and symbiont responses (B). Only terms explaining >1% of variance are displayed. Remaining unexplained variation (residuals) is indicated in light gray.

#### 1. Acropora pulchra

*Acropora pulchra* physiology exhibited significant variation across time, modulated by site, in both host (PERMANOVA P[time point:site]=0.033; PERMDISP P[site]=0.015; **Fig S3A**) and symbiont responses (PERMANOVA P[time point:site]=0.021; **Fig S4A**; **Table S9**; **Table S10**). However, time explained comparatively more variance (26.73%; RDA P=0.001; **Fig 6A**) and had a larger effect (Omega R2=0.39) relative to site (Omega R2=0.09; 8.02% variance explained, RDA P=0.001; **Table S10**; **Table S11**). All host physiological responses were affected by time (**Table S7**; **Fig S5**). Respiration rates were highest in January with antioxidant capacity, calcification, and protein content peaking in September, followed by elevated biomass in November (PC1 in **Fig S3D**; **Fig S5**). Site variation was observed in respiration rates (Linear mixed effect models (LMM) P[site]=0.002; **Fig S5D**) and protein content (LMM P[site]=0.003; **Fig S6C**), which were higher at the Orovau site, particularly during January-March (PC2 in **Fig S4D**).

*Acropora pulchra* symbiont physiology was explained to a similar extent by time (39.24%; RDA P=0.001; **Fig 6B**), with a small (2.46%), but significant (RDA P=0.004; **Table S11**), effect of site (PERMDISP P=0.002; **Table S10**). Time had a larger effect on multivariate physiology (Omega R2=0.25) than site (Omega R2=0.07; **Table S11**). All symbiont responses were affected by time (LMM P<0.01 for all metrics; PC1 in **Fig S4D**), except AQY (LMM P=0.051; **Table S8**). Symbiont density exhibited a clear peak in January-March (**Fig S6A**) while cell-specific chlorophyll and S:H biomass were highest in September (**Fig S6HI**).

#### 2. Pocillopora

Multivariate physiology in *Pocillopora* spp. varied across time, modulated by site, in both the host (PERMANOVA P[time:site]=0.001; **Fig S3B**) and symbiont (PERMANOVA P[time point:site]=0.003; **Fig S4B**; **Table S9**). Site explained a small, but significant, portion of host physiology (3.05%; RDA P[site]=0.002; **Fig 6A**). Time had a larger effect (Omega R2 = 0.24) than site (Omega R2 = 0.03). Host responses varied significantly between *P. tuahiniensis* and *P. meandrina* (PERMANOVA P=0.005; T**able S9**; **Table S10**; **Fig S7**), but with a small effect size (Omega R2=0.02).

All host responses except respiration (LMM P=0.281; **Fig S5D**) were significantly affected by time (LMM P<0.001 for all metrics; **Table S7**; **Fig S5**). Calcification (**Fig S5E**) peaked in March-September (PC2 in **Fig S3E**), biomass was highest in November (**Fig S5B**; PC1 in **Fig S3E**), and protein was lowest in November (**Fig S5C**; PC1 in **Fig S3E**). The effect of time on host biomass was modulated by site (LMM P=0.001; **Table S7**) with corals at Orovau retaining lower biomass than those at the other sites in November (**Fig S5B**). There was only a small effect of haplotype on host physiology in *P. meandrina* and *P. tuahiniensis* (Omega R2=0.02; **Fig S8**; **Table S12**).

Symbiont physiology was significantly explained by time (28.01%; RDA P=0.001; Omega R2 = 0.22; **Fig 6B**; **Table S11**) with all physiological responses affected (LMM P<0.01 for all metrics; **Table S8**). Symbiont responses varied significantly between holobionts (PERMANOVA P=0.001; PERMDISP P=0.481; **Table S10**; **Fig S7**) but effects were small (Omega R2=0.04; **Table S9**). Symbiont biomass (LMM P<0.001) and S:H biomass (LMM P<0.001) were higher in *P. meandrina* than *P. tuahiniensis* (**Fig S9**; **Table S13**). I_K_ exhibited a strong peak in March followed by a sharp decrease in September (**Fig S6E**) and AQY was elevated in both January and September (**Fig S6D**; PC1 in **Fig S4E**).

#### 3. Porites

Physiological responses in *Porites spp.* were significantly affected by site and time interactions in both the host (PERMANOVA P[time point:site]=0.012; **Fig S3C**) and symbiont (PERMANOVA P[time point:site]=0.001; **Fig S4C**; **Table S9**). In the host, time explained 27.98% of physiological variation (RDA P=0.001; Omega R2=-0.24; **Table S11**) with a small, but significant, amount of variation explained by site (1.69%, RDA P=0.001; Omega R2=0.04; **Fig 6A**). All host physiological responses were affected by time (LMM P<0.05; **Table S7**). Specifically, antioxidant capacity peaked in September (LMM P<0.001; **Fig S5A**) and respiration rates decreased across the time series (LMM P<0.001; **Fig S5D**). Host physiological metrics including antioxidant capacity, biomass, respiration rate, and calcification were higher in *Porites* spp. than *Pocillopora* spp. and *Acropora pulchra* (**Fig S5**). The effect of holobiont was significant in the host (PERMANOVA P=0.001; PERMDISP P=0.035; **Table S9**; **Table S10**; **Fig S10**) explaining 3.09% of variance (**Fig 6A**). Specifically, antioxidant capacity (LMM P<0.001; **Fig S11A**) and protein content (LMM P<0.001; **Fig S11C**) were higher in *P. evermanni* than *P. lobata/lutea* (**Table S12**). Combined site and holobiont effects (“Site&Holobiont” in **Fig 6**) are present due to the inability of the model to truly separate site and holobiont effects because *P. lobata/lutea* was not present at Matotia and holobiont effects were therefore confounded by site (**Fig 6**). Therefore, combined site and holobiont effects illustrate variance explained that cannot be attributed to either main effect individually.

Variation in symbiont physiological responses were significantly explained by time (18.12%; RDA P=0.001) and site (4.63%; RDA P=0.001; **Fig 6B**; **Table S11**). All symbiont responses except for symbiont biomass (LMM P=0.071) were influenced by time (LMM P<0.05; **Fig S6**; **Table S8**). P_MAX_ was higher in January-March (LMM P=0.003; **Fig S6C**), S:H biomass decreased in November (LMM P=0.001; **Fig S6I**), and AQY peaked in January and September (LMM P<0.001; **Fig S6D**; **Table S8**). The effect of holobiont on symbiont physiology was significant (PERMANOVA P=0.001; **Fig S10**) with holobiont and site:holobiont effects explaining 6.02% and 8.02%, respectively (RDA P=0.001; **Fig 6C**; **Table S11**). The effect of holobiont was equally as strong on symbiont physiology as time (Omega R2=0.16), which were both stronger than the effect of site (Omega R2=0.05; **Table S9**). Several symbiont physiological characteristics were higher in *P. evermanni,* including cell density (LMM P<0.001; **Fig S12A**), symbiont biomass (LMM P<0.001; **Fig S12B**), I_C_ (LMM P=0.002; **Fig S12F**), and symbiont:host biomass (LMM P<0.001; **Fig S12I**). Only cell-specific chlorophyll content was higher in *P. lobata/lutea* (LMM P=0.024; **Fig S12H**). Although the mean responses displayed in **Fig S6** visually show separation in responses by site, particularly at Matotia (e.g., **Fig S6B**), this is confounded by the differential presence of holobionts at Matotia and was therefore not statistically significant after accounting for holobiont as a random effect.

### C. Light and temperature changes across seasonality affect host and symbiont physiological responses

Physiological responses in the host and symbiont were significantly influenced by both light and temperature (RDA P<0.01; **Table S14**). Across all genera, light explained more variance in symbiont physiology than temperature, with the highest variance explained (34.3%) in *A. pulchra* (**Fig 7A**). Calcification and host protein content were positively correlated with light and increased during high light seasons, whereas biomass was negatively correlated and was higher in the cooler seasonal period (i.e., biomass in opposite direction from light arrows in **Fig 7BEH**). *Pocillopora* spp. symbiont physiology was also highly responsive to light (25.1%; **Fig 7D**) with *Porites* spp. symbiont physiology less responsive than the other two genera (13.1%; **Fig 7G**).

**Fig 7.**
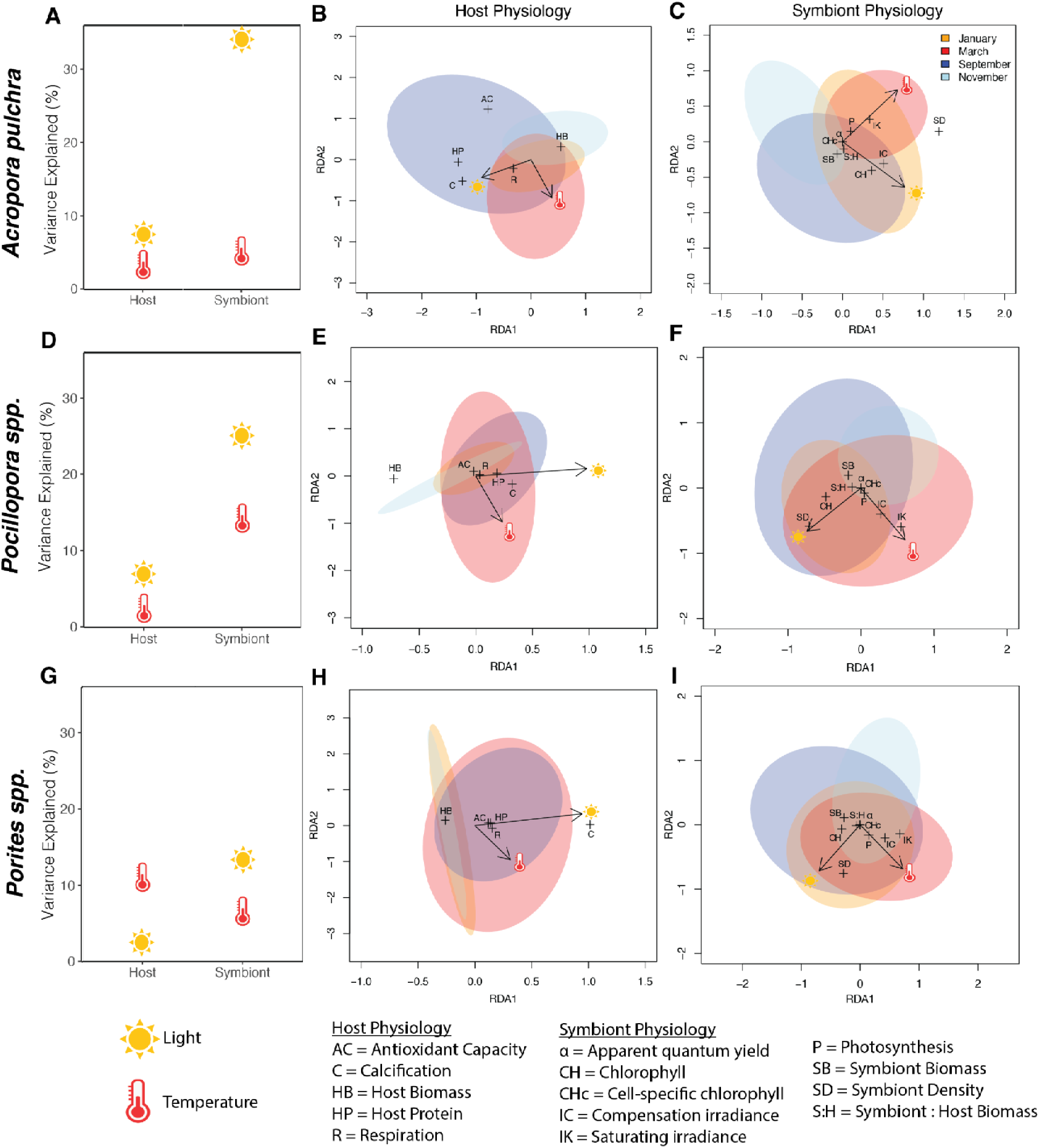
Variance partitioning analysis of temperature and light seasonal effects on physiology. Percent variance explained by light (sun icon) and temperature (thermometer icon) in multivariate physiology of the host and symbiont (x-axis) for each genus (A = *Acropora pulchra,* D = *Pocillopora spp.*, G = *Porites spp.*) as analyzed using redundancy analyses (RDA). Light and temperature effects on host and symbiont physiology were significant for all genera at P<0.05. RDA plots showing how light (sun icon and associated arrow) and temperature (thermometer icon with associated arrow) drive host (B, E, H) and symbiont (C, F, I) physiology for each genus. + indicates the location in the ordination and therefore association of each physiological metric with light and temperature drivers with metrics closer to the arrows being more strongly associated with the respective environmental variable. Metrics in opposite directions of variable arrows are negatively associated and metrics in the same direction as the arrows are positively associated. Metrics that are further from the plot origin have a stronger response to one or more environmental variables while those near the plot origin have weak associations. Longer environmental variable arrows indicate the strength of correlations with associated physiological metrics. Ellipses represent 95% confidence intervals (orange = January, red = March, dark blue = September, light blue = November).

Temperature explained the most variance in symbiont physiology in *Pocillopora* spp. (13.3%; **Fig 7D**) as compared to <10% variance explained in *Porites* spp. and *A. pulchra* (**Fig 7AG**), driven by positive correlations of photosynthetic parameters I_K_ and I_C_ with temperature (**Fig 7F**). Host physiology in all species was less responsive to light (2.4-7.5% variance explained; **Fig 7ADG**) with calcification being the primary light-driven response in the host (**Fig 7BEH**). Variance in *Porites* spp. host physiology was explained by temperature (10.0%) to a greater extent than *Pocillopora* spp. (1.6%) or *A. pulchra* (2.3%), which was driven by the positive relationship between temperature with respiration and a negative relationship between temperature and host biomass (**Fig 7H**).

### D. Variation in symbiont community composition is associated with variation in physiological responses in *Pocillopora* spp. and *Porites* spp

Variation in the ITS2 symbiont community composition of *A. pulchra* was not significantly related to either the host (dbRDA P=0.751) or symbiont physiology (dbRDA P=0.624; **Table S15**). However, variation in the symbiont community composition was related to the physiology of *Pocillopora* spp. and *Porites spp.* In *Pocillopora* spp., the symbiont community composition was significantly correlated with symbiont physiology (dbRDA P=0.001), but there was no significant relationship with host physiology (dbRDA P=0.886; **Table S15**). Symbiont metrics significantly related to community composition in *Pocillopora* spp. included symbiont biomass (dbRDA P=0.014), total chlorophyll (dbRDA P=0.001), cell-specific chlorophyll (dbRDA P=0.046), I_C_ (dbRDA P=0.024), and S:H biomass (dbRDA P=0.033; **Table S16**). The multivariate physiology of the symbiont constrained 14.09% of the variance in the symbiont community composition in *Pocillopora* spp. with the model explaining 7.42% (Adj. R^2^) of the variance (**Table S15**).

In contrast, variation in the symbiont community composition in *Porites* spp. was related to the physiology of both the host and the symbiont (dbRDA P[host]=0.001, P[symbiont]=0.001; **Table S15**). The multivariate physiology of the host constrained 14.17% of the variance in the symbiont community composition (model Adj. R^2^=10.84%, **Table S15)**, driven by significant correlations with host antioxidant capacity (dbRDA P=0.001) and biomass (dbRDA P=0.002; **Table S16**). The physiology of the symbiont constrained comparatively more of the variance in the symbiont community composition (22.58%, Adj. R^2^=17.01%, **Table S15**). Symbiont cell density (dbRDA P=0.002), symbiont biomass (dbRDA P=0.001), P_MAX_ (dbRDA P=0.038), and AQY (dbRDA P=0.002) were significantly correlated with symbiont community composition (**Table S16**).

## Discussion

Quantitative spatio-temporal host and symbiont physiology is key to both a mechanistic understanding of coral biology and improving the diagnostics of climate change consequences for reef building corals. Our study of seasonal acclimatization in three dominant Indo-Pacific reef-building coral genera reveals sharply contrasting physiological and symbiotic strategies that align with differences in environmental tolerance, which in turn shape their ecological distribution and success. The more environmentally sensitive and symbiotic generalist taxa, *Acropora* and *Pocillopora,* follow a symbiont-driven boom-and-bust cycle influenced primarily by changing light levels and, secondarily, by seasonal temperature fluctuations. In contrast, the environmentally resistant and symbiotic specialist, massive *Porites*, exhibits greater seasonal stability and greater host responsiveness to temperature, compared to light, across seasons. Further, cryptic lineages of *Pocillopora* and *Porites* exhibited distinct symbiont communities and thus physiological responses, highlighting the critical importance of identifying cryptic lineages and their symbiont communities to interpret physiology and performance in reef-building corals. Our findings support the hypothesis that the environmental sensitivity and ecological dynamics of *Acropora* and *Pocillopora* following disturbances can be driven by wide-ranging symbiotic fluctuations contributing to strain on the symbiosis (e.g., competition and destabilization (Cunning & Baker, 2013; McIlroy et al., 2020); excess reactive oxygen species (ROS)/reactive nitrogen species (RNS) production (Gibbin et al., 2014; Roth, 2014); nutrient stress (Cui et al., 2022; Krueger et al., 2020; Rädecker et al., 2021)), which increase the sensitivity to, and negative outcomes of, dysbiosis. Further, our work demonstrates the need for characterizing seasonal baselines of coral species for trait-based analysis (Edmunds & Putnam, 2020; Madin et al., 2016), stress tolerance assays (Cunning et al., 2024), and mechanistic interpretation (Helgoe et al., 2024; Scheufen, Krämer, et al., 2017), all of which are key to our capacity to forecast reef futures.

### A. Stable symbioses support year-round persistence in Porites compared to boom-and-bust dynamics in Acropora and Pocillopora

*Acropora* and *Pocillopora* are often categorized as competitive and weedy species (Darling et al., 2012), with diverse and flexible symbiotic relationships (Putnam et al., 2012), while in contrast, massive *Porites* display higher stress tolerance (Darling et al., 2012) with more stable, high-fidelity symbioses (Putnam et al., 2012). Our results clearly demonstrate divergent symbiotic seasonal acclimatization strategies in *Acropora, Pocillopora,* and *Porites,* representing a spectrum of generalist (high symbiotic seasonal acclimatization) to specialist (low symbiotic seasonal acclimatization) associations (i.e., higher vs lower variance explained in symbiont physiology by time in **Fig 6**). Previously widely documented ecological data, particularly following bleaching events, indicate generalist and specialist symbiotic and life history features have important implications for physiological stability and stress tolerance (Baird & Marshall, 2002; Hughes et al., 2018; Loya et al., 2001; McClanahan et al., 2004; Pratchett et al., 2013; Putnam et al., 2012; Silverstein et al., 2012). While these generalists or “boom-and-bust” strategies provide high acclimatization plasticity and flexibility, we posit that physiological flexibility can place strain on symbiotic nutritional relationships and therefore contribute to observed variation in holobiont ecological success (Loya et al., 2001). In particular, generalist relationships with high temporal variation in symbiont density and photophysiology (i.e., *Acropora*) may lead to increased symbiont competition, destabilization, and nutrient regulation imbalances that increase susceptibility to environmental stress, as compared to species with more stable symbiotic strategies (i.e., *Porites*).

In our seasonal study, the more environmentally sensitive taxa, *Acropora* and *Pocillopora*, hosted a wider diversity of symbiont taxa and displayed more dynamic “boom-and-bust” physiological responses, in comparison to the tolerant *Porites* corals that host specialized *Cladocopium* C15 lineages and exhibit more stable seasonal symbiont physiology. For example, *Porites* exhibited the greatest stability in symbiont and host biomass across seasonal periods, while *Acropora* and *Pocillopora* showed higher variability in photosynthetic rates, host biomass, symbiont biomass, and S:H biomass (**Fig S6**, **Fig S7**). Such symbiotic dynamics have holobiont consequences at a variety of levels. Specifically, symbiont density and performance have large consequences on the internal cellular physicochemical environment. For example, in a study of soft corals *Litophyton* sp. and *Rhytisma fulvum fulvum* from the Red Sea, increasing symbiont density led to decreased photosynthetic carbon translocation to the host and altered internal oxygen and carbon availability (Pupier et al., 2019). Symbiont taxa exhibit differing rates of photosynthesis (Brading et al., 2011, 2013) and/or carbon translocation (Allen-Waller & Barott, 2023; Stat et al., 2008), which creates varying and less predictable signals of pH, oxygen, and carbon to the host (Al-Horani, 2005; Dellaert & Putnam, 2023; Kühl et al., 1995). Seasonal fluctuations in symbiont community composition and physiological performance not only impact holobiont-level traits, but also impact the internal physiochemical environment that the coral host must tightly regulate.

Corals compartmentalize and maintain pH microenvironments that differ from the external environment. Earlier work with microprobes shows pH spatial variability across a polyp and colony (Al-Horani, 2005; Kühl et al., 1995), while more sophisticated imaging technologies, such as live confocal microscopy and pH-sensitive dye imaging, have provided quantification of intracellular pH (pHi; Barott et al., 2015; Venn et al., 2009). Coral cell pHi is regulated to an extent by the host (Venn et al., 2025) and buffered by the rates of photosynthesis of the symbionts (Gibbin et al., 2014; Laurent et al., 2013). Thus, variation in photosynthetic rates across symbiont species (Brading et al., 2011, 2013; Wall et al., 2020) and further influenced by symbiont density within the cells (Hoogenboom et al., 2010; Scheufen, Iglesias-Prieto, et al., 2017) creates internal environmental signals (Innis et al., 2021; Venn et al., 2025), requiring energetic investment from the host to maintain cellular homeostasis (Tresguerres et al., 2017). Therefore, in species with high temporal symbiont density and performance fluctuations (i.e., *Acropora* and *Pocillopora* in our study), the energetic demands of proton pumping (Capasso et al., 2021) and gene and protein regulation (Vidal-Dupiol et al., 2013) could place an energetic strain on the holobiont. Variation in photosynthetic rates also contributes to high oxygen flux and gradients across coral tissues (Kühl et al., 1995; Wangpraseurt et al., 2012, 2016). For example, microprobe work demonstrates that within the tissues of *Favia* and *Acropora*, oxygen can change >200% air saturation within a matter of minutes when switched from a dark to light condition (Kühl et al., 1995). Hypoxia and anoxia can drive the Hypoxia-Inducible Factor-mediated Hypoxia Response System (HIF-HRS) response in corals (Alderdice et al., 2021), with implications for metabolic reprogramming (Glass & Barott, 2025). Further, with higher photosynthetic oxygen production comes higher ROS generation and the potential increased need to ameliorate ROS-induced damage (Alderdice et al., 2022; Helgoe et al., 2024; Roth, 2014; Weis, 2008). Indeed, *Acropora* demonstrated the greatest variability in photosynthetic rates across seasons, with a 15-26% change between each seasonal time point (max. change in September), compared to 3-10% change in *Pocillopora* (max. change in November) and 3-20% change in *Porites* across seasons (max. change in September) (**Fig S7C**). Therefore, our findings that *Acropora* exhibited the largest temporal fluctuations in photosynthetic performance indicate that these corals may require disproportionately greater energetic allocation from the host to buffer against symbiont-driven variability in pH and oxygen dynamics, thereby placing the coral host in an energetically compromising position and contributing to environmental sensitivities.

While symbiotic flexibility may provide corals with the capacity to associate with symbiont taxa optimized for variable environmental conditions, flexibility in taxa associations may increase the risk of destabilization of symbiotic nutritional interactions. In *Acropora* particularly, large seasonal swings in metabolic rates and energy reserves (i.e., 16-17% change in host biomass, 9-40% change in symbiont biomass, and 11-39% change in respiration) may compromise host regulation of symbiont density (Cunning & Baker, 2013), by disrupting the balance and availability of key nutrients, particularly nitrogen, which plays a central role in host-mediated symbiont population control (Falkowski et al., 1993; Pogoreutz et al., 2017; Radecker et al., 2015; Rädecker et al., 2023; Xiang et al., 2020). Therefore, in periods of highly fluctuating metabolism and physiological state, nutrient limitation and symbiont regulation may become less predictable, allowing symbiont communities to proliferate (Falkowski et al., 1993; McIlroy et al., 2022; Radecker et al., 2015; Rädecker et al., 2023). In turn, growing symbiont populations may increase nutrient demands, leading to nitrogen and phosphorus limitation (Morris et al., 2019) and reduced efficiency of carbon translocation to the host and energy deficits (Tremblay et al., 2014). *Acropora* was the only taxa to exhibit symbiont growth during the seasonal warm period of March (14% increase), while *Pocillopora* (11% reduction) and *Porites* (5% reduction) showed slight decreases in density during this period. Periods of symbiont overpopulation and destabilization are associated with increased bleaching risk (Cunning & Baker, 2013; Wooldridge, 2020). Dysbiosis due to metabolic imbalance is further exacerbated by the production of excess ROS and RNS that can damage both symbiont and host, triggering oxidative stress pathways that compromise symbiotic stability and increase bleaching (Gibbin et al., 2014; Roth, 2014). Indeed, in our study, we observed that antioxidant capacity was greatest in *Porites* and lowest in *Acropora* and *Pocillopora* during January-March, periods when symbiont densities were at their peak, with these effects most pronounced in *Acropora*. Furthermore, *P. lutea* at Wuzhizhou Island has high fidelity associations with C15 and exhibits lower symbiont densities than more flexible species (i.e., *Acropora* and *Montipora*) and was more tolerant under anthropogenic disturbances (Chen et al., 2024). This suggests that taxa with more flexible symbiotic strategies must dedicate more energy towards oxidative stress management in periods of symbiont growth. We hypothesize that high seasonal variability in symbiotic associations in taxa such as *Acropora* and *Pocillopora* contributes to increased bleaching susceptibility in these species, in contrast to more seasonally stable *Porites*.

Given the variable conditions and the demands of hosting multiple types of symbionts, there may be increased symbiont competition in generalist species (i.e., *Acropora* and *Pocillopora*). Such dynamic population and metabolic changes can contribute to a symbiotic situation of competition for nutrient resources (McIlroy et al., 2020). Critically, uptake and translocation of carbon and nitrogen shift depending on symbiont density and host:symbiont ratios (McIlroy et al., 2022). Nutrient limitation shapes the exchange of nutritional resources between host and symbiont and is a core component of symbiotic stability and productivity (Cui et al., 2023; Ezzat et al., 2015; Rädecker et al., 2023; Shantz & Burkepile, 2014). Nitrogen (Ezzat et al., 2015; Krueger et al., 2020; Radecker et al., 2015; Rädecker et al., 2021) and phosphorus (J. Li et al., 2023; Rosset et al., 2017; Wiedenmann et al., 2013) availability is regulated to control symbiont growth and optimize carbon translocation from symbiont to host. However, when symbiont densities surge, as seen in *Acropora* during peak growth periods in January-March in this study (i.e., 14% increase in March followed by a 24% decrease in September and 50% decrease in November relative to January), nutrient demand may outpace supply, leading to nutrient imbalance and limitation (Cui et al., 2022; Falkowski et al., 1993; Krueger et al., 2020; Xiang et al., 2020). Under these conditions, symbionts may retain a greater portion of photosynthetically-produced carbon, reducing translocation to the host and therefore availability of carbon in the holobiont (Cunning et al., 2017; Rädecker et al., 2018; Wooldridge, 2013), with excessive symbiont densities showing reduced carbon translocation (Anthony et al., 2009; Hoogenboom et al., 2010; Wooldridge, 2020). Reductions in carbon sharing may create energetic shortfalls in the host that may be detrimental if occurring during or before periods of high metabolic demand, such as reproduction and bleaching (Leinbach et al., 2021). In contrast, *Porites* exhibited more stable symbiont densities during seasonal warm periods (4-6% reductions in March-September) and high-fidelity associations with C15 symbionts and therefore may experience more consistent nutrient dynamics and a steady supply of carbon throughout seasonal cycles, allowing this taxa to maintain energy reserves. Indeed, we documented stable host biomass throughout seasonal cycles in *Porites* with stable host biomass and biomass gains (5-17% increases in biomass across all seasons) compared to losses in September in *Acropora* and *Pocillopora* (16% and 11% loss, respectively). This biomass stability may allow *Porites to* avoid seasonal energy store losses following periods of high metabolic demands, as seen in more flexible taxa, *Pocillopora* and *Acropora*. Over time and repeated seasonal cycles, higher stability in nutrient and carbon exchange dynamics and biomass storage may contribute to the observed higher stress tolerance across studies in *Porites* (Baird & Marshall, 2002; Hughes et al., 2018; Loya et al., 2001; McClanahan et al., 2004; Pratchett et al., 2013).

### B. Identifying seasonal baselines and physiological strategies is critical for interpreting ecological outcomes

Our findings suggest that symbiont competition in generalist hosts, while allowing for potentially beneficial physiological flexibility, may introduce instability that compromises host tolerance under fluctuating or stressful conditions. Stability in symbiont densities and ratios may facilitate more consistent energy acquisition throughout seasonal periods in *Porites,* providing the energy required for consistent performance during environmental disturbances. In environments that are within tolerance ranges for specialized host-symbiont associations, taxa like *Porites* may outperform due to greater consistency in energy production and acquisition throughout seasonal cycles. Overall, our study suggests that both symbiotic strategy and host life history contribute to the stability of coral responses to environmental variation. Corals with fast growth and flexible, “boom-and-bust” dynamics, such as *Acropora*, which exhibit higher metabolic and symbiotic flexibility, may achieve short-term gains but may also be prone to sharp declines, as documented during bleaching events (Baird & Marshall, 2002; Hughes et al., 2018; Loya et al., 2001; McClanahan et al., 2004; Pratchett et al., 2013). In contrast, corals like *Porites*, which maintain slower growth and more stable, specialized symbiotic associations, appear to sustain more consistent energy supply across seasons. This stability may reduce the risk of symbiotic stress under environmental extremes and avoid the energetic volatility inherent to boom-and-bust strategies. Thus, while flexibility can offer short-term advantages, long-term resilience may depend more on the steadiness of stable partnerships and conservative growth strategies.

Our study further demonstrates that it is critical to characterize the seasonal physiological baseline of corals to understand and interpret performance across space and time. We found that both symbiont performance (e.g., chlorophyll content, symbiont density, photosynthetic rate) and host energetic reserves (e.g., biomass and metabolic rates) exhibited strong seasonal variation. These shifts varied in magnitude and pattern across taxa, reflecting taxa-specific metabolic strategies for coping with environmental variation. For example, *Porites* corals maintained relatively stable host energy reserves while *Pocillopora* and *Acropora* followed symbiont-driven boom-and-bust fluctuations in symbiont and host metrics. Therefore, seasonal taxa-specific context is essential for interpreting changes in performance and physiology. Because of this variation, we recommend that researchers provide seasonal physiological baselines to contextualize measurements of performance and stress tolerance. Previous studies have shown that thermal tolerance thresholds can shift seasonally, likely reflecting changes in metabolic state. For example, *P. verrucosa* corals in the Red Sea show up to a 3°C variation in thermal thresholds depending on the season with *Acropora* corals showing a 1°C variation (García et al., 2024). Periods of reduced tolerance were seen in cooler seasonal periods and may be a result of shifts in metabolism between seasons (García et al., 2024). Further, *P. damicornis* winter bleaching thresholds were 1°C lower than summer thresholds (Berkelmans & Willis, 1999). In contrast, juvenile *P. damicornis* corals reared in seasonal cool temperatures exhibited greater biomass reserves and higher thermal tolerance (Huffmyer et al., 2021). Our results underscore these prior findings and provide detailed physiological evidence that seasonal metabolic baselines are not only variable but are likely drivers of resilience and performance.

### C. Identifying the presence of cryptic lineages is critical for interpreting physiological responses

Identifying cryptic coral species is critical for interpreting physiological responses because each species hosts distinct symbiont communities and exhibits unique physiological traits. Cryptic species are common in *Pocillopora* (Burgess et al., 2021), and symbiont composition is tightly linked to host species identity (Johnston et al., 2022; Turnham et al., 2021). In our study, *Pocillopora tuahiniensis* hosted *Cladocopium pacificum,* whereas *P. meandrina* hosted *Cladocopium latusorum* (Johnston et al., 2022; Millán-Márquez et al., 2024), and these communities did not vary by site. These species-specific associations were accompanied by differences in symbiont traits including chlorophyll content and biomass, indicating that differences in symbiont communities can lead to distinct physiological responses across cryptic holobiont lineages. *Porites* showed high fidelity to *Cladocopium* C15 symbionts in our study, yet each lineage (i.e., *P. evermanni* and *P. lobata/lutea*) hosted distinct C15 subtypes, similar to observations in *Porites* corals in Palau (Grupstra et al., 2024) and Kiritimati (Starko et al., 2023). These differences were also seen in function with symbiont (e.g., symbiont density and photosynthetic traits) and host physiology (e.g., biomass and antioxidant capacity) varying with community composition. Indeed, other work in *Porites* demonstrated variation in photophysiology and thermal tolerance between host lineages, as well as variation in associated bacterial communities (Grupstra et al., 2024).

Collectively, our results underscore that physiological responses cannot be generalized across morphologically similar coral species within the same genus. Cryptic species differ in both their symbiotic relationships (Turnham et al., 2021) and their physiological performance (Burgess et al., 2021), making species-level resolution essential for understanding coral environmental resistance and resilience. In this study, the *Porites lobata/lutea* lineage was only found at two of our sites (Vaipahu and Orovau), while *Porites evermanni* was found at all three sites. Because physiological responses varied between *Porites* species, without species identification we would have falsely attributed variation in physiology between species to site effects. This has also been identified as a potential confounding factor in other studies (Starko et al., 2023). Cryptic species in *Pocillopora* and *Porites* corals are difficult to identify visually with morphology alone (Forsman et al., 2009; Grupstra et al., 2024; Johnston et al., 2022), and we recommend the inclusion of molecular identification of coral genera and associated symbionts to more accurately account for cryptic lineage effects.

## Conclusions

In this study, we describe physiological strategies for seasonal acclimatization in three major reef-building coral genera in Mo’orea, French Polynesia ranging from the stable, high-fidelity strategy in *Porites*, to the boom-and-bust pattern of *Acropora*, and the muted but flexible response of *Pocillopora*. These distinct physiological strategies may influence resilience to environmental stress and reef trajectories under changing conditions and highlight the importance of considering both host and symbiont contributions to holobiont function. We emphasize that characterizing seasonal baselines is critical for understanding coral performance and making meaningful predictions about tolerance to future stress. Together, our findings underscore that physiological and symbiotic strategies play important roles in environmental tolerance that shape ecological distribution and success.

## Acknowledgements

As guests, we honor and acknowledge the Mā’ohi community, we give thanks for the land and water resources of Polynesia and the traditional custodians of the land on which this experimental work was conducted on the island of Mo’orea. Māuruuru roa. With respect to the spelling of Tahitian words, we endeavored to follow the Te Fare Vāna□a, transcription system that is adhered to by a large segment of the Tahitian community but also recognize other community members follow the Raapoto transcription system where the island name of Mo’orea is, for example, spelled without the ‘eta (i.e., Mo’orea).

## Author Contributions

Hollie M. Putnam, Roger M. Nisbet, Ross Cunning, Jose M. Eirin-Lopez, and Steven B. Roberts conceived the ideas and design of the project and acquired funding. Hollie M. Putnam, Ross Cunning, Jose M. Eirin-Lopez, Dennis Conetta, Emma L. Strand, Juliet M. Wong, Kevin H. Wong, Danielle M. Becker, Kristina X. Terpis, Francis J. Oliaro, and Zoe Dellaert collected data. Ariana S. Huffmyer, Ferdinand Pfab, Emma L. Strand, Serena Hackerott, and Hollie M. Putnam analyzed the data. Ariana S. Huffmyer, Emma L. Strand, Serena Hackerott, and Hollie M. Putnam wrote the original manuscript and Ariana S. Huffmyer, Emma L. Strand, and Hollie M. Putnam revised the manuscript. All authors approved the final version of the manuscript.

## Data Availability Statement

All data and code are openly available on GitHub (https://github.com/urol-e5/timeseries) and as a static release (https://github.com/urol-e5/timeseries/releases/tag/v1.0) and will additionally be deposited to a permanent data repository (Zenodo) following review.

