## Supporting Information for "Seasonal plasticity of symbiotic strategies clarifies coral holobiont resistance and resilience"

### Supporting Figures

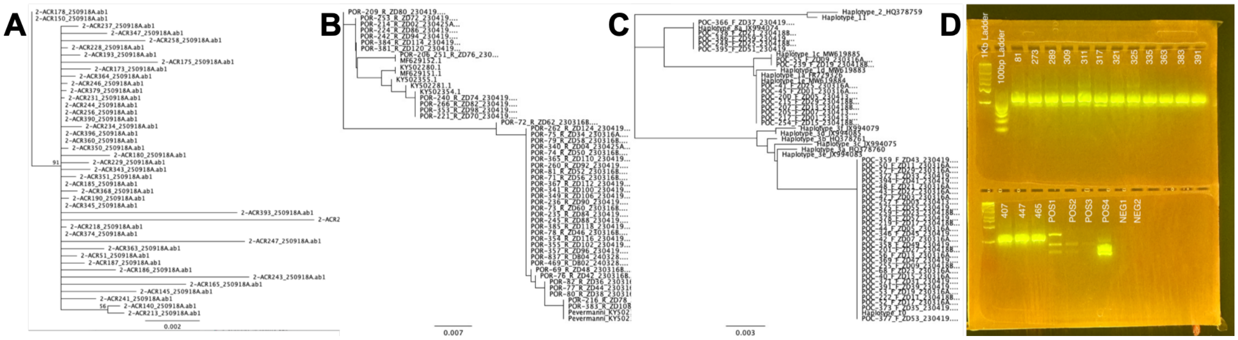

**Fig S1**. Species trees for A) *Acropora*, B) *Porites*, and C) *Pocillopora* (mtORF) and D) RFLP gel for *Pocillopora* (PocHistone).

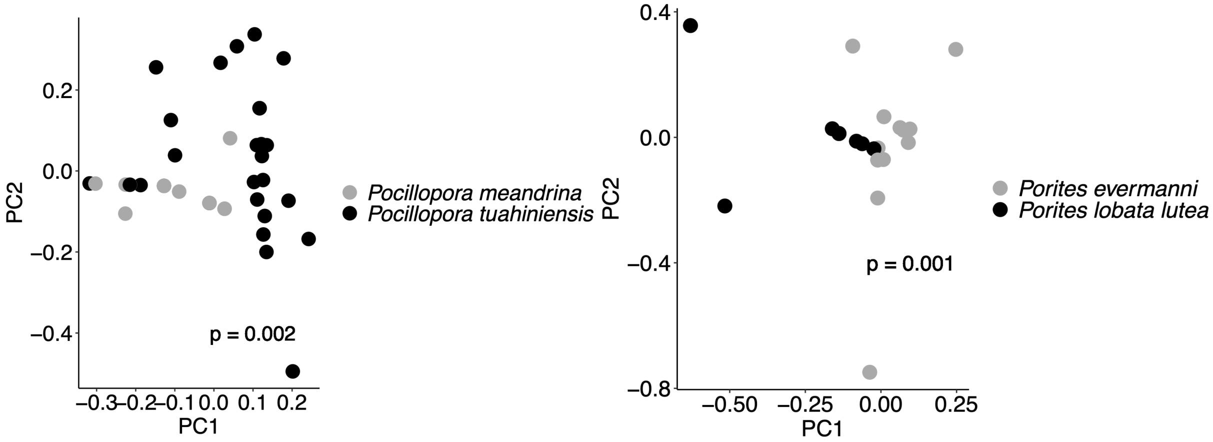

**Fig S2. Principal components analysis of symbiont community.** Symbiont communities visualized by holobiont identity for *Pocillopora* (left) and *Porites* (right) genera. In *Pocillopora*, gray indicates *Pocillopora meandrina* and black indicates *Pocillopora tuahiniensis*. In *Porites*, gray indicates *Porites evermanni* and black indicates *Porites lobata lutea*. P-values indicate significance of permutational analysis of variance (PERMANOVA) tests of the effect of holobiont on symbiont community composition (ITS2 type profile relative abundance).

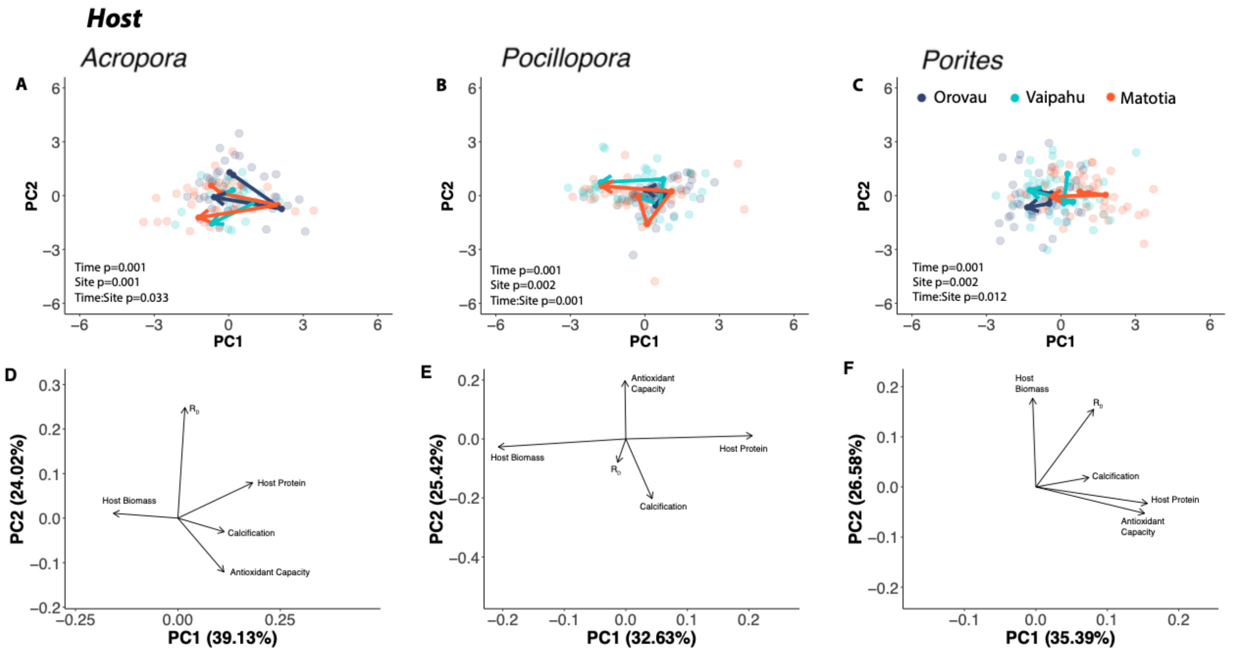

**Fig S3. Host physiological responses across sites and time points.** Multivariate physiological trajectories in (A) *Acropora pulchra*, (B) *Pocillopora* spp., and (C) *Porites* spp. across site and time as visualized with principal components analyses. Trajectory arrows display the centroid of multivariate physiology of each species across time points, with the arrows beginning at the centroid of January 2020 samples and ending at the centroid of November 2020 samples. Color indicates site (Orovau = blue, Vaipahu = cyan, Matotia = orange). Biplots displayed for *Acropora pulchra* (D), *Pocillopora* spp. (E), and *Porites* spp. (F). Text indicates significance of site, time point, and the interaction using permutational analysis of variance tests. $R_D$ indicates respiration.

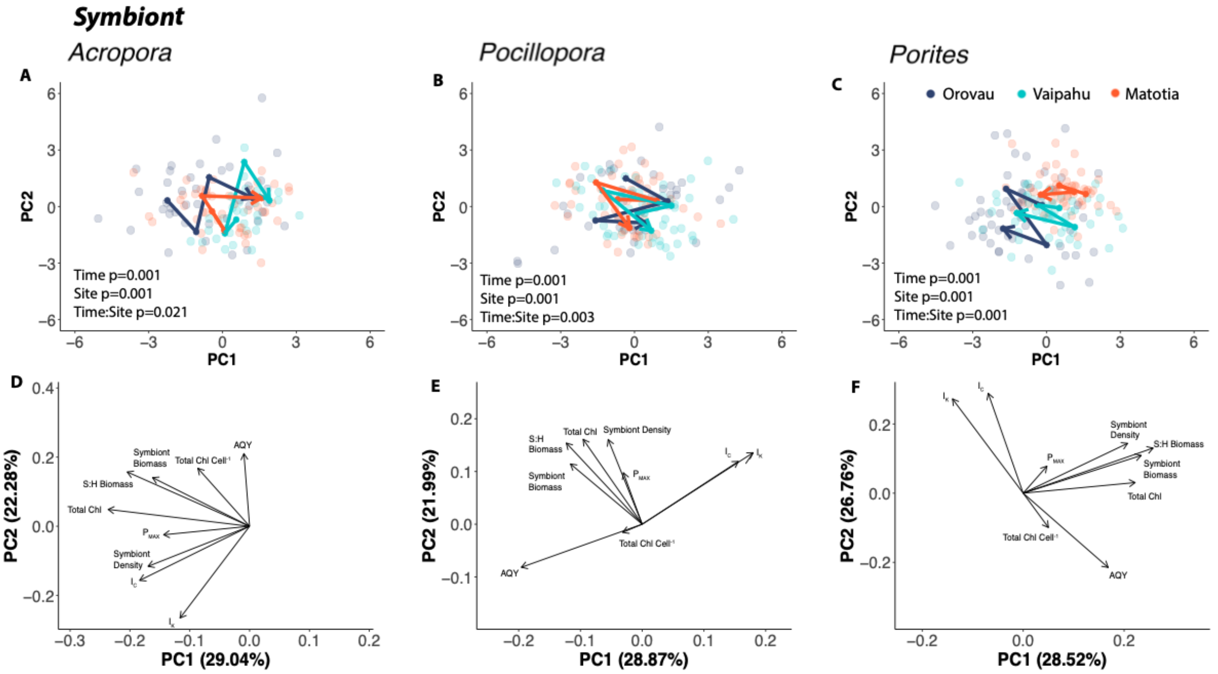

**Fig S4. Symbiont physiological responses across sites and time points.** Multivariate physiological trajectories in (A) *Acropora pulchra*, (B) *Pocillopora* spp., and (C) *Porites* spp. across site and time as visualized with principal components analyses. Trajectory arrows display the centroid of multivariate physiology of each species across time points, with the arrows beginning at the centroid of January 2020 samples and ending at the centroid of November 2020 samples. Color indicates site (Orovau = blue, Vaipahu = cyan, Matotia = orange). Biplots displayed for *Acropora pulchra* (D), *Pocillopora spp.* (E), and *Porites spp.* (F). Text indicates significance of site, time point, and the interaction using permutational analysis of variance tests. $P_{MAX}$ indicates maximal photosynthesis; $I_C$ indicates compensation irradiance; $I_K$ indicates saturating irradiance.

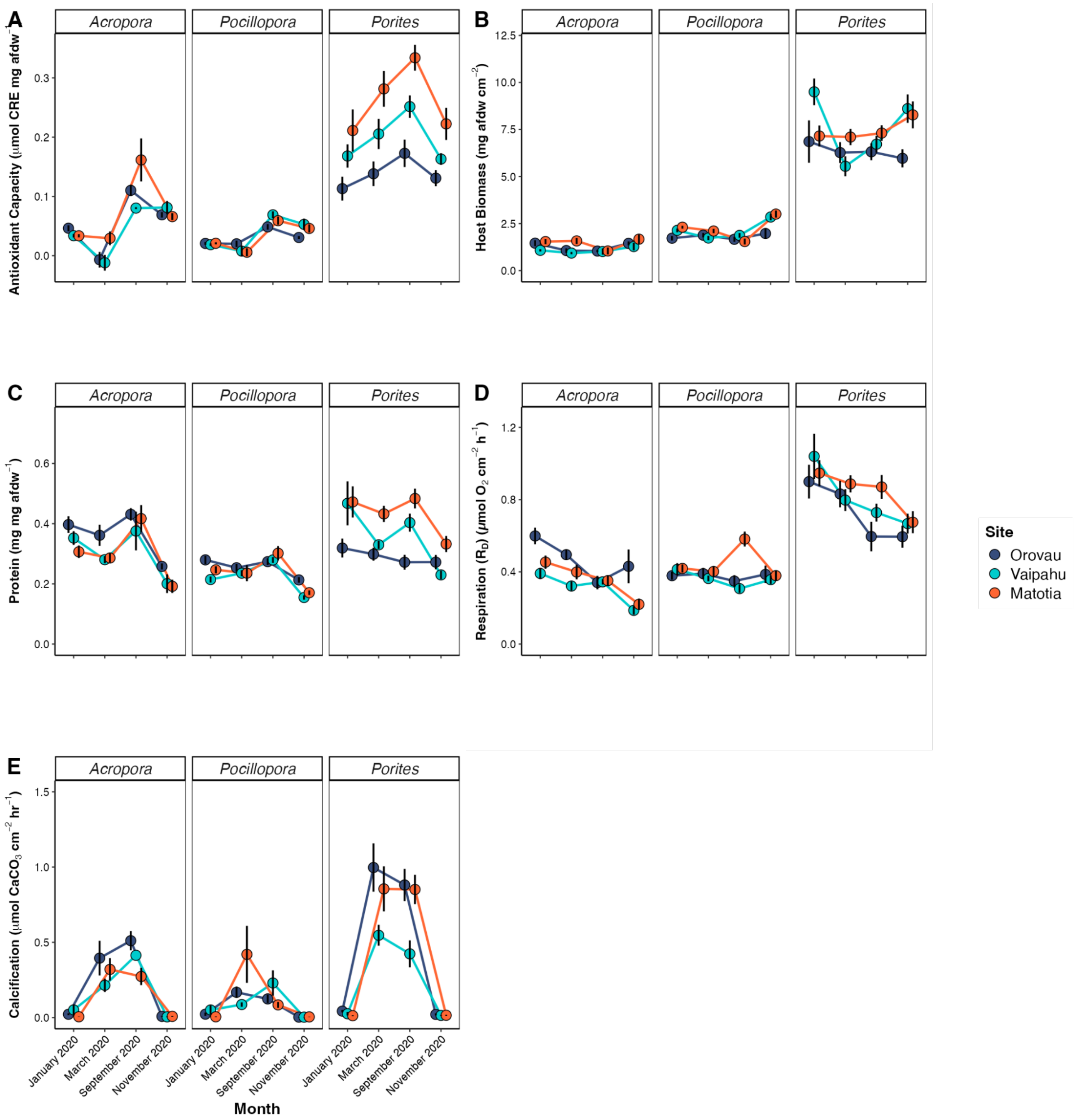

**Fig S5. Host responses across site and time point for each genus.** Mean ± standard error of mean of host responses across time point (x-axis) and site (blue=Orovau, cyan=Vaipahu, orange=Matotia). Plots are faceted by genus. Host responses include (A) antioxidant capacity (μmol copper reducing elements mg $AFDW^{-1}$), (B) host biomass (mg AFDW $cm^{-2}$), (C) host protein (mg protein mg $AFDW^{-1}$), (D) respiration ($R_D$; μmol $O_2$ $cm^{-2}$ $h^{-1}$), (E) calcification (μmol $CaCO_3$ $cm^{-2}$ $h^{-1}$). Time points are ordered as January, March, September, and November 2020.

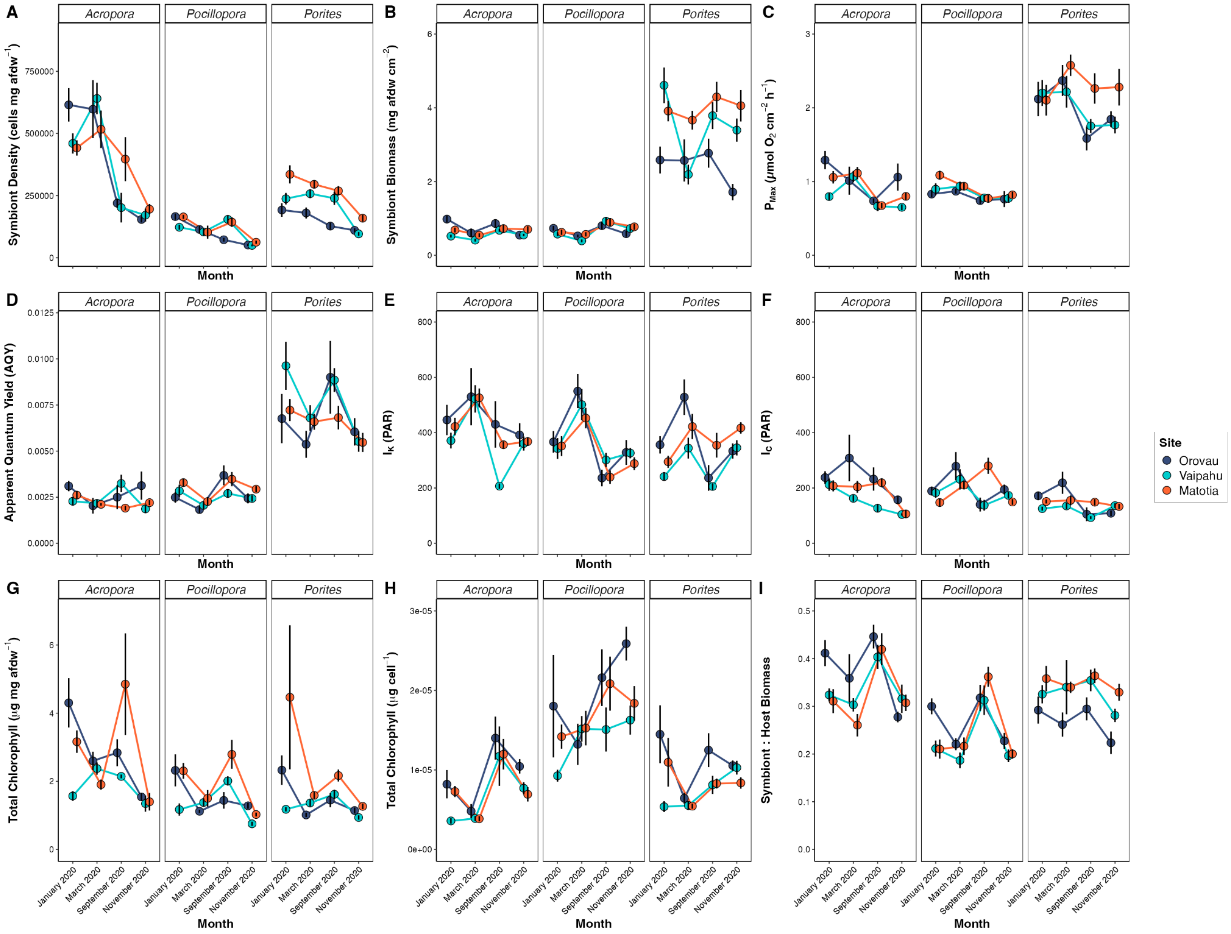

**Fig S6. Symbiont responses across site and time point for each genus.** Mean ± standard error of mean of symbiont responses across time point (x-axis) and site (blue=Orovau, cyan=Vaipahu, orange=Matotia). Plots are faceted by genus. Responses include (A) symbiont cell density (cells mg $AFDW^{-1}$), (B) symbiont biomass (mg AFDW $cm^{-2}$), (C) maximal photosynthesis ($P_{MAX}$; μmol $O_2$ $cm^{-2}$ $h^{-1}$), (D) apparent quantum yield (AQY; expressed as a proportion), (E) saturating irradiance ($I_K$; PAR), (F) compensation irradiance ($I_C$; PAR), (G) total chlorophyll (chl *a* + chl $c_2$) (μg pigment mg $AFDW^{-1}$), (H) total cell-specific chlorophyll (chl *a* + chl $c_2$) (μg pigment $cell^{-1}$), (I) S:H biomass. Time points are ordered as January, March, September, and November 2020.

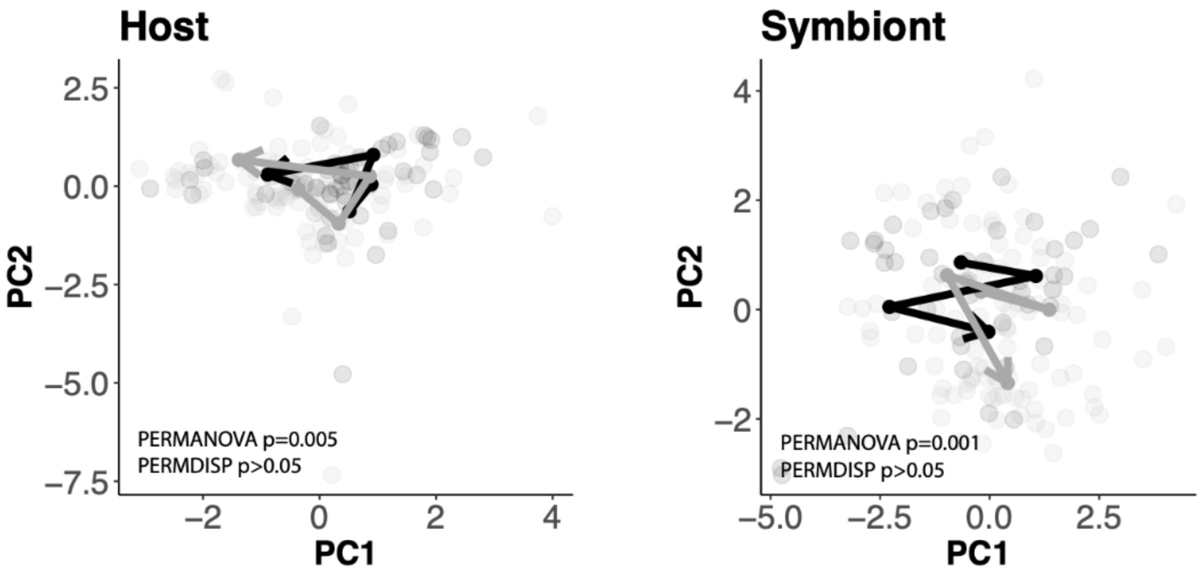

**FIg S7.** Physiological responses across time points for each *Pocillopora* holobiont - *P. meandrina* (black) and *P. tuahiniensis* (gray) in host (left) and symbiont (right) responses. Trajectory arrows display the centroid of multivariate physiology of each holobiont across time points, with the arrows beginning at the centroid of January 2020 and ending at the centroid of November 2020. P-values indicate significance of permutational analysis of variance (PERMANOVA) and permutational analysis of dispersion (PERMDISP) tests.

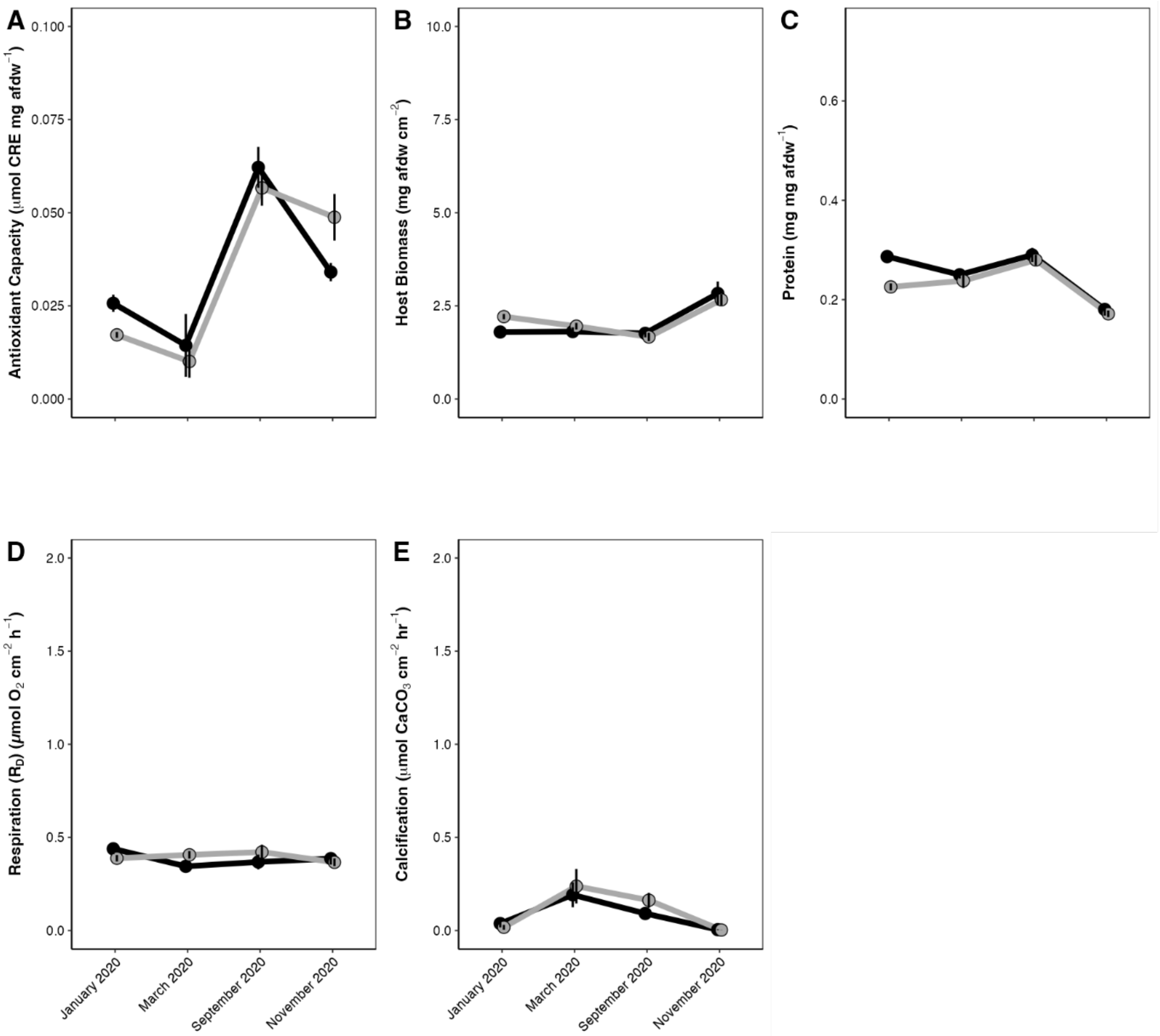

**Fig S8. Host responses across site and time point for each haplotype in the *Pocillopora* genus.** Mean ± standard error of mean of host responses across time point (x-axis) and haplotypes in *Pocillopora* (black=*P. meandrina*, gray=*P. tuahiniensis*). Host responses include (A) antioxidant capacity (μmol CRE mg $AFDW^{-1}$), (B) host biomass (mg AFDW $cm^{-2}$), (C) host protein (mg protein $AFDW^{-1}$), (D) respiration ($R_D$; μmol $O_2$ $cm^{-2}$ $h^{-1}$), (E) calcification (μmol $CaCO_3$ $cm^{-2}$ $h^{-1}$). Time points are ordered as January, March, September, and November 2020.

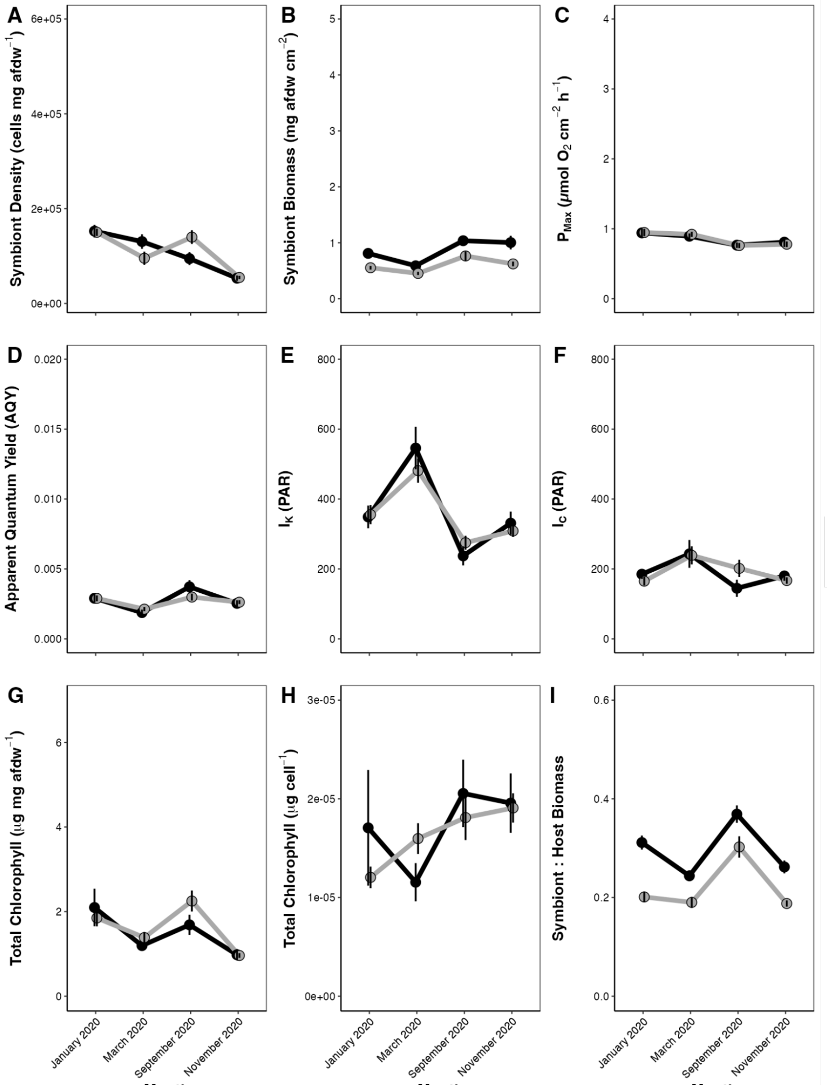

**FIg S9. Symbiont responses across site and time point for each holobiont in the *Pocillopora* genus.** Mean ± standard error of mean of symbiont responses across time point (x-axis) and haplotypes in *Pocillopora* (black=*P. meandrina*, gray=*P. tuahiniensis*). Host responses include (A) symbiont cell density (cells mg AFDW$^{-1}$), (B) symbiont biomass (mg AFDW cm$^{-2}$), (C) maximal photosynthesis ($P_{MAX}$; µmol $O_2$ cm$^{-2}$ h$^{-1}$), (D) apparent quantum yield (AQY; expressed as a proportion), (E) saturating irradiance ($I_K$; PAR), (F) compensation irradiance ($I_C$; PAR), (G) total chlorophyll (chl *a* + chl $c_2$) (µg pigment mg AFDW$^{-1}$), (H) total cell-specific chlorophyll (chl *a* + chl $c_2$; µg pigment cell$^{-1}$), (I) S:H biomass. Time points are ordered as January, March, September, and November 2020.

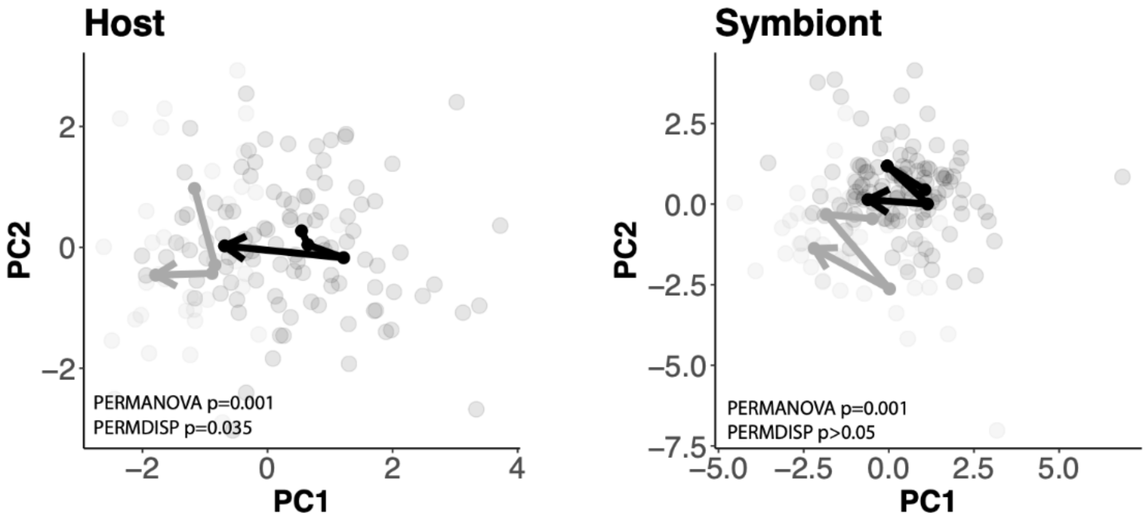

**Fig S10.** Physiological responses across time points for each *Porites* holobiont - *P. evermanni* (black) and *P. lobata/lutea* (gray) for host (left) and symbiont (right) responses. Trajectory arrows display the centroid of multivariate physiology of each holobiont across time points, with the arrows beginning at the centroid of January 2020 and ending at the centroid of November 2020. P-values indicate significance of permutational analysis of variance (PERMANOVA) and permutational analysis of dispersion (PERMDISP) tests.

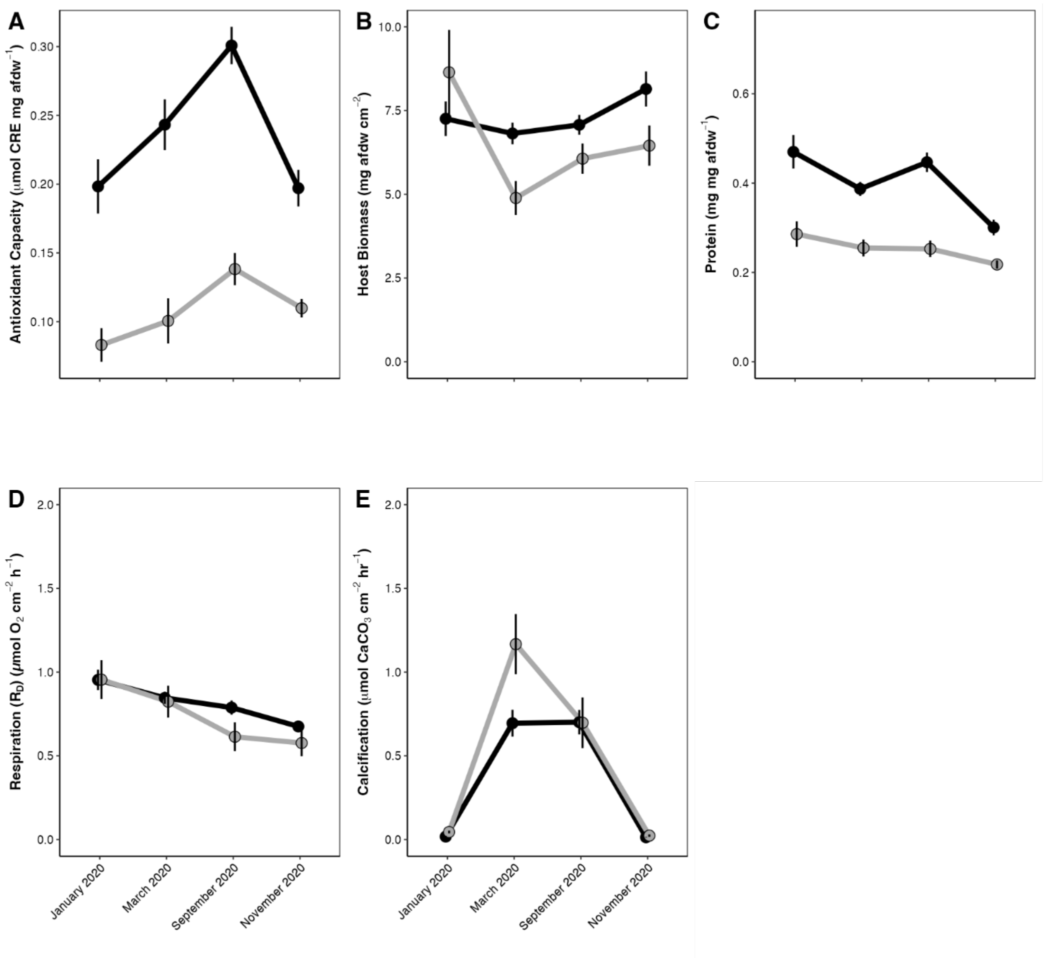

**Fig S11. Host responses across site and time point for each holobiont in the *Porites* genus.** Mean ± standard error of mean of host responses across time point (x-axis) and haplotypes in *Porites* (black=*P. evermanni*, gray=*P. lobata/lutea*). Host responses include (A) antioxidant capacity (μmol CRE mg AFDW$^{-1}$), (B) host biomass (mg AFDW cm$^{-2}$), (C) host protein (mg protein AFDW$^{-1}$), (D) respiration ($R_D$; μmol $O_2$ cm$^{-2}$ h$^{-1}$), (E) calcification (μmol $CaCO_3$ cm$^{-2}$ h$^{-1}$). Time points are ordered as January, March, September, and November 2020.

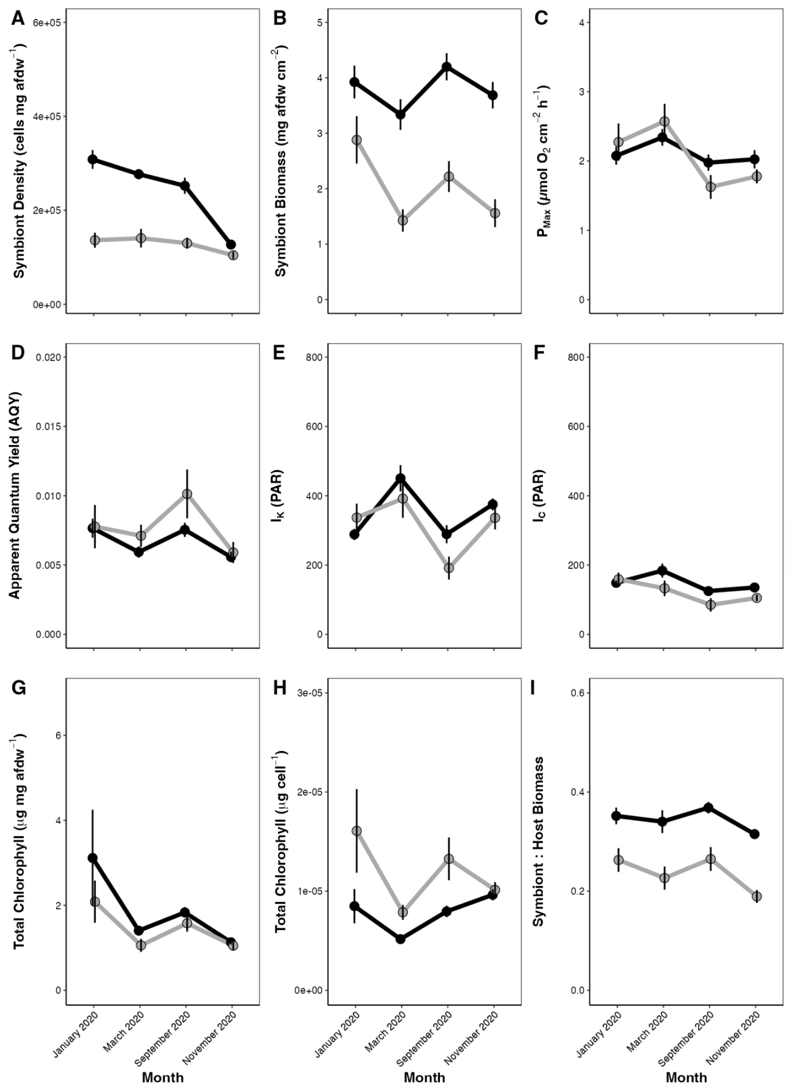

**Fig S12. Symbiont responses across site and time point for each holobiont in the *Porites* genus.** Mean ± standard error of mean of symbiont responses across time point (x-axis) and and haplotypes in *Porites* (black=*P. evermanni*, gray=*P. lobata/lutea*). Host responses include (A) symbiont cell density (cells mg AFDW$^{-1}$), (B) symbiont biomass (mg AFDW cm$^{-2}$), (C) maximal photosynthesis ($P_{MAX}$; μmol $O_2$ cm$^{-2}$ h$^{-1}$), (D) apparent quantum yield (AQY; expressed as a proportion), (E) saturating irradiance ($I_K$; PAR), (F) compensation irradiance ($I_C$; PAR), (G) total chlorophyll (chl *a* + chl $c_2$) (μg pigment mg AFDW$^{-1}$), (H) total cell-specific chlorophyll (chl *a* + chl $c_2$; μg pigment cell$^{-1}$), (I) S:H biomass. Time points are ordered as January, March, September, and November 2020.

### Supporting Methods

A. Photosynthesis-irradiance (PI) curves

Following acclimation, each fragment was placed into individual acrylic respiration chambers (620 mL) with a magnetic stir bar and aerated filtered seawater (FSW; 100 µm). Each photosynthesis-irradiance (PI) curve run included eight coral fragments and two chambers with filtered seawater (FSW) as blanks. Chambers were placed on magnetic stir plates and held in a water bath maintained at ~27°C with a recirculating pump and heater (Finnex 300W Titanium Heater, Burnaby, British Columbia, Canada). Temperature (PreSens Pt1000) and fiber-optic oxygen probes (Presens dipping probe; DP-PSt7-10-L2.5-ST10-YOP) were inserted into each chamber. Chambers were exposed to ten increasing light levels for 10 min each (0 [dark], 18, 68, 113, 169, 243, 499, 709, 844, and 1025 µmol photons $m^2$ $s^1$; PAR) with LED aquarium lighting (Prime 16HD Reef Aquarium Lights, AquaIllumination). Oxygen concentration (µmol $L^{-1}$) and temperature (°C) were measured every 1 sec. An underwater cosine corrected sensor (MQ-510 Quantum Meter, spectral range of 389–692 ± 5 nm, Apogee Instruments, Logan, UT, USA) was used to measure light levels at each chamber position prior to each measurement.

Oxygen consumption (under dark conditions) and production (under light conditions) rates were calculated for each light level interval using localized linear regressions (alpha=0.2, percentile rank method) in the *LoLinR* package (Olito et al., 2017) on selected data points from oxygen measurements at 20 sec intervals and then normalized to chamber volume to produce µmol $O_2$ $sec^{-1}$. Mean oxygen rates were calculated for blank chambers and subtracted from coral oxygen rates. Rates were then normalized to surface area and calculated as µmol $O_2$ $cm^{-2}$ $h^{-1}$.

Oxygen rates (µmol $O_2$ $cm^{-2}$ $h^{-1}$) from each light level (PAR) interval were then used to generate PI curves using the quadratic equation using flexible start values (Equation 1) (Falkowski & Raven, 2013).

*Equation (1)*

$$Oxygen\ (\mu mol\ O_2\ cm^{-2}\ h^{-1}) = P_{Max} \times \frac{AQY \times PAR}{\sqrt{{P_{Max}}^2 + (AQY \times PAR)^2}} - R_D$$

where $P_{Max}$ is the maximal photosynthetic rate, $AQY$ is the apparent quantum yield, $PAR$ is the light value, and $R_D$ is dark respiration. Model estimates of $P_{Max}$ (µmol $O_2$ $cm^{-2}$ $h^{-1}$), $AQY$, and $R_D$(µmol $O_2$ $cm^{-2}$ $h^{-1}$) were extracted for each colony at each time point. In addition, saturating irradiance $I_K$ (light value (PAR) at which initial slope crosses $P_{Max}$) for each colony was calculated as in Equation 2.

*Equation (2)*

$$I_K = \frac{P_{Max}}{AQY}$$

Compensation irradiance $I_C$ (light value (PAR) at which photosynthesis equals respiration) was also calculated for each colony as in Equation 3.

*Equation (3)*

$$I_C = \frac{P_{Max} \ x \ R_D}{AQY \ x \sqrt{{P_{Max}}^2 - {R_D}^2}}$$

B. Instantaneous calcification rates

After each PI curve assay, the oxygen and temperature probes were removed from the acrylic respiration chambers and each individual chamber was immediately filled and sealed with FSW to measure instantaneous calcification using the total alkalinity (TA) anomaly technique (Chisholm & Gattuso, 1991). Prior to incubations, duplicate initial 125 mL water samples were collected from the incoming water source (details on sampling below). Following a 90-minute incubation period at 28°C and 500 PAR (above saturating irradiance, see **Fig S7E**), each coral fragment was immediately frozen in liquid nitrogen and stored at -40 °C for additional physiological analyses. Water samples (125 mL) were collected from each coral (*n*=8) and blank (*n*=2) chamber following each incubation run. Salinity, pH, and temperature measurements were collected for each initial and final water sample using a Thermo Scientific Orion Star A222 Conductivity Portable Meter (Thermo Fisher Scientific, Franklin, MA, USA), InLab Expert Pro pH Electrode (Mettler Toledo, Port Melbourne, VIC, Australia), and a Digital Traceable Thermometer (Control Company 5-077-8, accuracy=0.05°C, resolution=0.001°C, Webster, TX, USA). Initial and final water samples were preserved with 75 µL of 50% saturated mercuric chloride ($HgCl_2$) solution. Total alkalinity was measured by following the open cell potentiometric titration (SOP3b; Dickson et al., 2007) procedure using a Mettler Toledo Titrator Excellence T5 (#30252672), Rondolino (#51108500), pH probe (DGi115-SC), and hydrochloric acid (Dickson Lab Batch A1) at the University of Rhode Island. Prior to each titration, the pH probe was calibrated with a three-point calibration using pH 4.0, 7.0, and 10.0 solutions. A Certified Reference Material (CRM; Dickson Lab Batch #180) sample was run as a standard to confirm <0.01% error. Calcification rates were calculated following the total alkalinity (TA) anomaly technique (Chisholm & Gattuso, 1991). TA values were normalized to salinity and corrected for change in TA in the blank samples. TA was normalized to surface area and incubation time to calculate calcification rates as µmol $CaCO_3$ $cm^{-2}$ $hr^{-2}$.

C. Surface area

After the coral tissue was separated from the skeleton using the airbrushing method above, the coral skeletons were placed in a drying oven (Fisher Scientific Isotemp Oven) at 60°C for at least 4 h until dry. Pre-weighed wooden dowels of known dimensions were used to determine a standard curve of mass change of wax-dipped dowels against geometrically calculated surface area ($R^2$>0.9; Stimson & Kinzie, 1991). The surface area of the dried coral fragments was measured by first weighing the dried coral skeleton and then dipping fragments once into a 65°C Minerva paraffin wax bath (Monroe, GA, USA) for 2 sec and then quickly rotating the coral skeleton in the air at a standardized rate (10 revolutions over 2 sec). Coral skeletons were cooled for 10 min before their final mass (g) was measured. The calibration curve was then used to determine the surface area of each fragment ($cm^2$) (Stimson & Kinzie, 1991; Veal et al., 2010).

D. Tissue biomass

Biomass was measured on both the symbiont and host tissue fractions as ash-free dry weight (AFDW). 5 mL of homogenate was centrifuged at 3,500 x g $min^{-1}$ for 3 min (Fisher Scientific accuSpin 3R), resulting in 4 mL of supernatant (host fraction) and an algal pellet (symbiont fraction), which was resuspended in 1 mL of cold 1X PBS. The host and symbiont fractions were aliquoted into pre-burned (450°C for 4-6 h) aluminum pans and were dried in a drying oven at 80°C for 24 h. Once fully dried, weight of pans was recorded and then placed in a muffle furnace at 450°C for 4-6 h. AFDW was calculated as the difference between dry weight and burned weight normalized to sample volume and fragment surface area to generate mg AFDW $cm^{-2}$.

E. Host soluble protein and total antioxidant capacity (TAC)

To calculate host soluble protein and total antioxidant capacity (TAC), a 1 mL host tissue homogenate aliquot was thawed on ice and centrifuged at 10,000 rpm at 4°C for 10 min (Eppendorf Centrifuge 5415D). The supernatant was removed and the sample was briefly vortexed. Protein content was measured using the Thermo Scientific Pierce BCA Protein Assay Kit with 25 µL of each sample according to the manufacturer's instructions. Protein content was calculated as absorbance at 562 nm against a diluted albumin (BSA) standard curve and normalized to homogenate volume and biomass to calculate soluble host protein concentration (mg protein mg $AFDW^{-1}$). Tissue physiological metrics were normalized to tissue biomass (Edmunds & Gates, 2002).

Antioxidant capacity (TAC) was measured with 20 µL of host tissue used in the Cell BioLabs OxiSelect TAC Assay Kit according to the manufacturer's instructions. Antioxidant capacity (mM uric acid equivalents; UAE) was calculated as the absorbance at 490 nm against a uric acid standard curve. TAC was calculated as µmol Copper Reducing Equivalents (CRE) per mg AFDW (µmol CRE mg $AFDW^{-1}$) as in Equation 4.

*Equation (4)*

$$CRE\ \frac{umol}{mg\ AFDW} = \frac{UAE\ (mM)\ \times\ 2189\ \ \mu M\ Cu++/\ mM\ uric\ acid\ \times\ Homogenate\ volume\ (L)}{biomass\ (mg)}$$

F. Chlorophyll content

To measure chlorophyll *a* and $c_2$ pigment concentration, the symbiont pellet was thawed at room temperature. Pigments were extracted by adding 1 mL of 100% acetone to symbiont pellets, vortexed, and then placed in a dark fridge at 4°C for 24 h. Samples were then centrifuged for 3 min at 16,000 rcf (Eppendorf Centrifuge 5415D). 200 µL of the supernatant containing extracted pigments were transferred to a 96-well quartz microplate (Hellma® Analytics, USA), and the absorbance of each sample was measured at 630, 663, and 750 nm on a plate spectrophotometer (Synergy HTX Multi-Mode Reader, BioTek, USA). Chlorophyll *a* (Equation 5a) and $c_2$ concentrations (Equation 5b) were calculated from the respective equations specific for dinoflagellates in 100% acetone (Eqn 1ab; (Jeffrey & Humphrey, 1975) and corrected for path length (0.66 cm; Hellma Analytics Quartz Microplate; Markham, Ontario Canada) (Equation 5). Chlorophyll *a* and chlorophyll $c_2$ content were normalized to biomass (µg pigment mg $AFDW^{-1}$) and symbiont cell density (described below; µg pigment $cell^{-1}$).

*Equation (5a)*

$$Chl\ a = \frac{(11.43\ x\ (E_{663\ nm}\ -\ E_{750\ nm})\ -\ 0.64\ \times\ (E_{630\ nm}\ -\ E_{750\ nm}))}{PATH\ LENGTH\ (0.66)}$$

*Equation (5b)*

$$Chl\ c_2\ = \frac{(27.09\ x\ (E_{630\ nm}\ -\ E_{750\ nm})\ -\ 3.63\ \times\ (E_{633\ nm}\ -\ E_{750\ nm}))}{PATH\ LENGTH\ (0.66)}$$

G. Molecular identification

We conducted genetic identification of host haplotypes in *Pocillopora* spp. and *Porites* spp. (Burgess et al., 2021; Forsman et al., 2009). *Pocillopora* species were identified by amplifying the mitochondrial open reading frame (mtORF) region as described by (Burgess et al., 2021; Johnston et al., 2018) using primers from (Flot et al., 2008): FatP6.1 (5′-TTTGGGSATTCGTTTAGCAG-3′) and RORF (5′-SCCAATATGTTAAACASCATGTCA-3′). We then used the PocHistone 3 region to distinguish species in the mtORF haplotype 1a (*P. meandrina* and *P. eydouxi*) as described (Johnston et al., 2018) using the primers PocHistoneF (5′-ATTCAGTCTCACTCACTCACTCAC-3′) and PocHistoneR (5′-TATCTTCGAACAGACCCACCAAAT-3′). PCR master mixes (25 µL total reaction volume) contained 12.55 µL of EmeraldAmp GT PCR Master Mix (TaKaRa Bio USA Inc. Cat # RR310B), 0.32 µL of forward and reverse primers at 10 µM, 1 µL of template DNA, and 10.80 µL of nuclease free water (Invitrogen UltraPure CAT # 10977015). We included positive controls as DNA from previously successfully amplified samples and a negative control with water in place of template DNA. The mtORF region was amplified using a PCR protocol of a single denaturation set of 94°C for 60 sec followed by 30 cycles of 94°C for 30 sec for denaturation, 53°C for 30 sec for annealing, and 72°C for 75 sec for extension and a final incubation of 72°C for 5 min. The PocHistone region was amplified using the same PCR protocol with the exception of the extension step conducted at 72°C for 60 sec. PCR products were assessed with a 1.5% agarose gel in TAE for 30 min at 80 V.

*Porites* species were identified using the coral nuclear histone region spanning H2A to H4 (i.e., H2) (Tisthammer et al., 2020) using the primers: zH2AH4f (5′-GTGTACTTGGCTGCYGTRCT-3′) and zH4Fr (5′-GACAACCGAGAATGTCCGGT-3′). This marker can delineate *Porites evermanni* apart from the clade containing *P. lobata/lutea*. The coral nuclear histone region was amplified using a PCR protocol of a single denaturation set of 94°C for 2 min followed by 34 cycles of 96°C for 20 sec for denaturation, 58.5°C for 20 sec for annealing, and 72°C for 90 sec for extension and a final incubation of 72°C for 5 min. Products were assessed on a 1.5% agarose gel in TAE for 30 min at 80 V to confirm only one band of approx. 1500 bp was recovered.

*Acropora* samples were identified by amplifying the mitochondrial control region using primers from (Vollmer & Palumbi, 2002): CRf (5'-GCTTAGACAGGTTGGTTGATTGCCC-3') and CO3r (5'-CTCCCAAATACATAATTTGAACTAA-3'). The mitochondrial control region was amplified using a PCR protocol of a single denaturation set of 95°C for 3 min followed by 35 cycles of 94°C for 30 sec for denaturation, 53°C for 30 sec for annealing, and 72°C for 60 sec for extension and a final incubation of 72°C for 5 min.

PCR products for all species were cleaned using ethanol precipitation, with 1/10th of the volume of 3 M sodium acetate (Fisher Cat. AAJ61928AE), followed by an addition of 3 times the total volume of the mixture of cold 100% ethanol and overnight incubation at -20°C. DNA was precipitated by centrifugation at 15,000 rcf for 30 min at room temperature and the pellet was then washed with 70% ethanol twice, dried, and resuspended in 30 µL of 1 M Tris-HCl, pH 8.0 (Fisher Cat. 15568025). DNA was assessed with dsDNA Qubit (Broad Range) and Nanodrop. Sanger sequencing using the same primers as described above was performed at the URI Genomics and Sequencing center using Applied Biosystems BigDye Terminator v3.1. Sequences were aligned and analyzed using Geneious Alignment in GENEIOUS PRIME 2020.2.4 as described in (Harnay & Putnam, 2023). Pairwise alignment of reads was completed using Clustal Omega (Sievers et al., 2011) and neighbor-joining trees were constructed (Zhang & Sun, 2008) with Jukes-Cantor genetic distance model (Jukes & Cantor, 1969) for both forward and reverse reads.

For species identification, Pocillopora mtORF was compared against the following Haplotype 1a FR729326 (Flot et al., 2010); Haplotype 1c MW619885, Haplotype 1d MW619883, Haplotype 1e MW619884 (Burgess et al., 2021); Haplotype 2 HQ378759, Haplotype 3a HQ378760, Haplotype 3b HQ378761 (Pinzón & LaJeunesse, 2011); Haplotype 3c JX994075, Haplotype 3d JX994085, Haplotype 3e JX994083, Haplotype 3f JX994079, Haplotype 3h JX994072, Haplotype 5a JX994073, Haplotype 8a JX994074 (Pinzón et al., 2013);

Haplotype 10 OP418359 (Johnston & Burgess, 2023); Haplotype 11 KF381328 Forsman et al 2013 (KF381328) (Johnston et al., 2022) full sequence. When *Pocillopora* species were identified as Haplotype 1 with mtORF, RFLP was conducted according to (Johnston et al., 2018). For *Porites* identification, H2 sequences were compared to sequences from (Brown et al., 2020) *P. evermanni* (KY502351, KY502352) and *P. lobata/lutea* (MF629151, MF629152, KY502280, KY502281, KY502354, KY502355). For *Acropora* identification, no reference canonical A*. pulchra* sequence is available from Moorea, so we examined all of the sequences from this study only to test the hypothesis that we had only one group of sequences. Forward and reverse sequences were too short to be joined, so alignment and tree building was conducted on 600 bp of the reverse sequences, where the total sequence number available for analysis was highest.

H. Symbiont ITS2 identification

Thermal cycling consisted of an initial denaturation at 98°C for 2 min, followed by 35 cycles of 98°C for 10 sec, 56°C for 30 sec, and 72°C for 30 sec, and a final extension step at 72°C for 5 min following (Hume et al., 2018). Each 25 µL reaction contained 12.5 µL Q5 MasterMix (New England Biolabs), a unique combination of barcoded forward and reverse primers at 5 µM, and 2 µL template DNA. PCR products were purified and normalized to a concentration of ~1 ng $\mu L^{-1}$ using Charm Just-a-plate 96 PCR Normalization and Purification Kit (Charm Biotech) and measured in triplicate on Qubit™ 3.0 fluorometer using the 1X dsDNA High Sensitivity kit (Invitrogen). Pooled libraries were denatured and diluted following the standard Illumina sequencing protocol and sequenced on an Illumina MiSeq using a 500-cycle v2 reagent kit (250 bp paired-end reads) and custom sequencing primers to initiate forward, reverse and index reads (Kozich et al., 2013).

**Table S1. Sample sizes of the number of colonies sampled across time points for each genus, holobiont, and site.** Holobiont indicates host haplotype and associated symbionts.

| Genus | Holobiont | Site | Number of colonies measured per time point | | | |
|---|---|---|---|---|---|---|
| | | | January | March | September | November |
| *Acropora* | *Acropora pulchra* | Vaipahu | 13 | 9 | 2 | 6 |
| | | Orovau | 14 | 6 | 10 | 12 |
| | | Matotia | 15 | 9 | 9 | 11 |
| *Pocillopora* | *Pocillopora meandrina* | Vaipahu | 3 | 3 | 3 | 3 |
| | | Orovau | 8 | 9 | 7 | 7 |
| | | Matotia | 3 | 3 | 2 | 2 |
| *Pocillopora* | *Pocillopora tuahinensis* | Vaipahu | 12 | 11 | 10 | 12 |
| | | Orovau | 5 | 6 | 6 | 5 |
| | | Matotia | 12 | 12 | 10 | 11 |
| *Porites* | *Porites evermanni* | Vaipahu | 12 | 11 | 12 | 11 |
| | | Orovau | 6 | 6 | 4 | 4 |
| | | Matotia | 15 | 15 | 13 | 12 |
| *Porites* | *Porites lobata lutea* | Vaipahu | 3 | 3 | 3 | 3 |
| | | Orovau | 9 | 9 | 8 | 8 |
| | | Matotia | 0 | 0 | 0 | 0 |

**Table S2. Effect of genus on multivariate physiology.** Permutational analysis of variance (PERMANOVA) and permutational analysis of dispersion (PERMDISP) analyses conducted for biological level (combined responses, host, symbiont) separately with genus as the main effect. Bold indicates P<0.05. DF = degrees of freedom; SS = sum of squares. Tests run with 999 permutations.

| Test | Biological Level | Main Effect | DF | SS | R2 | F value | P-value |
|---|---|---|---|---|---|---|---|
| PERMANOVA | Combined | genus | 2 | 1872.90 | 0.36 | 104.00 | **0.001** |
| | Host | genus | 2 | 835.20 | 0.43 | 145.30 | **0.001** |
| | Symbiont | genus | 2 | 1174.50 | 0.32 | 95.20 | **0.001** |
| PERMDISP | Combined | genus | 2 | 1.03E+12 | | 70.87 | **<0.001** |
| | Host | genus | 2 | 164.28 | | 98.49 | **<0.001** |
| | Symbiont | genus | 2 | 1.21E+12 | | 82.91 | **<0.001** |

**Table S3. Pairwise multivariate analysis comparisons between holobionts within the *Pocillopora* and *Porites* genera.** Permutational analysis of variance **(**PERMANOVA) and permutational analysis of dispersion (PERMDISP) analyses conducted for biological level (combined responses, host, symbiont) separately. Posthoc comparisons between holobionts within each genus conducted using pairwise PERMANOVA and Tukey HSD comparisons (PERMDISP tests). Bold indicates P<0.05. DF = degrees of freedom; SS = sum of squares. Holobiont indicates host haplotype and associated symbionts.

| **Genus** | **Comparison** | **Biological Level** | **Pairwise PERMANOVA** | | | | **Pairwise PERMDISP** | |
|---|---|---|---|---|---|---|---|---|
| | | | DF | SS | F | Adjusted P-value | Difference | Adjusted P-value |
| *Pocillopora* | *P. tuahiniensis - P. meandrina* | Combined | 1 | 1.25E+12 | 0.362 | 0.551 | 1272.123 | 0.999 |
| | | Host | 1 | 1.543 | 3.136 | 0.690 | 0.077 | 0.989 |
| | | Symbiont | 1 | 3.15E+06 | 8.46E-04 | 1.000 | 4184.311 | 0.999 |
| *Porites* | *P. lobata/lutea - P. evermanni* | Combined | 1 | 3.24E+11 | 49.733 | **0.001** | -40077.049 | 0.102 |
| | | Host | 1 | 18.037 | 3.357 | 0.590 | 0.396 | 0.163 |
| | | Symbiont | 1 | 3.732 | 6.05E+01 | **0.010** | -35248.049 | 0.148 |

**Table S4. Permutational multivariate analysis of symbiont communities within each genus.** PERMANOVA analyses conducted on relative abundance of ITS2 profiles for each genus separately. Holobiont, time point, site, and the interaction of time and site were included as main effects. Bold indicates P<0.05. DF = degrees of freedom; SS = sum of squares. Holobiont refers to the host genetic haplotype and associated symbiont communities.

| Genus | Main Effect | DF | SS | R2 | F value | P-value |
|---|---|---|---|---|---|---|
| *Acropora* | site | 2 | 0.19 | 0.01 | 0.44 | 0.77 |
| | time point | 3 | 0.30 | 0.01 | 0.47 | 0.853 |
| | site:time point | 6 | 0.38 | 0.02 | 0.30 | 0.986 |
| | residuals | 107 | 22.64 | 0.96 | | |
| *Pocillopora* | holobiont | 1 | 14.28 | 0.12 | 20.31 | **0.002** |
| | time point | 2 | 2.88 | 0.02 | 2.05 | 0.052 |
| | site:time point | 3 | 0.98 | 0.01 | 0.46 | 0.955 |
| | holobiont | 6 | 2.70 | 0.02 | 0.64 | 0.913 |
| | residuals | 145 | 102.00 | 0.83 | | |
| *Porites* | holobiont | 1 | 31.06 | 0.29 | 63.04 | **0.001** |
| | site | 2 | 1.99 | 0.02 | 2.02 | 0.060 |
| | time point | 3 | 0.60 | 0.01 | 0.40 | 0.986 |
| | site:time point | 6 | 1.10 | 0.01 | 0.37 | 0.998 |
| | residuals | 149 | 73.408 | 0.68 | | |

**Table S5**. Analysis of variance (3-way ANOVA) tests for effects of site, ITS2 profile, and holobiont on relative abundance of Symbiodiniaceae taxa that comprise >1% of total relative abundance. DF = degrees of freedom; SS = sum of squares. Bold indicates P<0.05.

| Genus | Main Effect | DF | SS | F | P-value |
|---|---|---|---|---|---|
| | site | 2 | 0.00 | 0.00 | 1.000 |
| ***Acropora*** | profile | 4 | 24.29 | 147.11 | **<0.001** |
| | site:profile | 6 | 0.08 | 0.24 | 0.961 |
| | site | 2 | 0.00 | 0.00 | 1.000 |
| | profile | 13 | 7.06 | 15.18 | **<0.001** |
| | holobiont | 1 | 0.00 | 0.00 | 1.000 |
| ***Pocillopora*** | site:profile | 26 | 1.66 | 1.78 | **0.011** |
| | site:holobiont | 2 | 0.00 | 0.00 | 1.000 |
| | profile:holobiont | 13 | 3.31 | 7.12 | **<0.001** |
| | site:profile:holobiont | 26 | 0.64 | 0.69 | 0.876 |
| | site | 2 | 0.00 | 0.00 | 1.000 |
| | profile | 20 | 12.78 | 37.42 | **<0.001** |
| | holobiont | 1 | 0.00 | 0.00 | 1.000 |
| ***Porites*** | site:profile | 40 | 3.24 | 4.74 | **<0.001** |
| | site:holobiont | 1 | 0.00 | 0.00 | 1.000 |
| | profile:holobiont | 20 | 5.83 | 17.07 | **<0.001** |
| | site:profile:holobiont | 20 | 0.14 | 0.42 | 0.988 |

**Table S6. Variance explained by main effects of multivariate physiology.** Variance partitoning analyses conducted for each genus, biological level with effects of site, time, and holobiont identity for any term explaining >1% of variance.The percent of the variance explained by each factor individually. F test statistics and P-values for each main effect determined from ANOVA-like permutation analyses of partial redundancy analyses for each individual main effect controlling for all other main effects.

| Genus | Biological Level | Effect | Variance Explained (%) |
|---|---|---|---|
| *Acropora* | Combined | time point | 38.37 |
| | | site | 3.13 |
| | | residuals | 59.64 |
| | Host | time point | 26.73 |
| | | site | 8.02 |
| | | residuals | 66.67 |
| | Symbiont | time point | 39.24 |
| | | site | 2.46 |
| | | residuals | 59.39 |
| *Pocillopora* | Combined | time point | 28.40 |
| | | site | 1.33 |
| | | residuals | 70.40 |
| | Host | time point | 18.28 |
| | | site | 3.05 |
| | | site:holobiont | 1.26 |
| | | residuals | 77.48 |
| | Symbiont | time point | 28.01 |
| | | residuals | 71.64 |
| *Porites* | Combined | time point | 19.33 |
| | | site | 3.86 |
| | | holobiont | 5.55 |
| | | site:holobiont | 6.96 |
| | | residuals | 63.81 |
| | Host | time point | 27.98 |
| | | site | 1.69 |
| | | site:holobiont | 3.09 |
| | | residuals | 67.74 |
| | Symbiont | time point | 18.12 |
| | | site | 4.63 |
| | | holobiont | 6.02 |
| | | site:holobiont | 8.02 |
| | | residuals | 62.04 |

**Table S7. Univariate linear mixed effect model analysis of host responses.** All responses were log+1 transformed. Linear mixed effect models were conducted for each genus and included time point, site, and their interactions as main effects with colony nested within holobiont (for *Porites* and *Pocillopora* only) as a random intercept. Significance determined with Type III analysis of variance tests. SS = sum of squares; Num DF = numerator degrees of freedom; Den DF = denominator degrees of freedom. Bold indicates P<0.05. CRE indicates copper reducing elements. AFDW indicates ash-free dry weight.

| Response | Genus | Main Effect | SS | Num DF | Den DF | F value | P-value |
|---|---|---|---|---|---|---|---|
| Antioxidant Capacity (µmol CRE mg AFDW-1) | *Acropora* | time point | 0.092 | 3 | 79.09 | 34.72 | **<0.001** |
| | | site | 0.004 | 2 | 58.35 | 2.81 | 0.068 |
| | | time point:site | 0.014 | 6 | 78.21 | 2.72 | **0.019** |
| | *Pocillopora* | time point | 0.043 | 3 | 123.00 | 33.43 | **<0.001** |
| | | site | 0.001 | 2 | 123.00 | 1.06 | 0.349 |
| | | time point:site | 0.005 | 6 | 123.00 | 2.00 | 0.071 |
| | *Porites* | time point | 0.119 | 3 | 94.65 | 21.51 | **<0.001** |
| | | site | 0.013 | 2 | 43.25 | 3.55 | **0.037** |
| | | time point:site | 0.019 | 6 | 94.59 | 1.74 | 0.119 |
| Host Soluble Protein (mg protein mg AFDW-1) | *Acropora* | time point | 0.231 | 3 | 75.41 | 27.23 | **<0.001** |
| | | site | 0.038 | 2 | 46.22 | 6.66 | **0.003** |
| | | time point:site | 0.011 | 6 | 74.31 | 0.66 | 0.680 |
| | *Pocillopora* | time point | 0.111 | 3 | 121.23 | 30.04 | **<0.001** |
| | | site | 0.012 | 2 | 114.96 | 4.69 | **0.011** |
| | | time point:site | 0.015 | 6 | 121.17 | 2.09 | 0.059 |
| | *Porites* | time point | 0.180 | 3 | 97.76 | 17.81 | **<0.001** |
| | | site | 0.024 | 2 | 24.98 | 3.49 | **0.039** |
| | | time point:site | 0.077 | 6 | 97.66 | 3.79 | **0.002** |
| Host Biomass (mg AFDW cm-2) | *Acropora* | time point | 0.323 | 3 | 77.49 | 3.14 | **0.030** |
| | | site | 0.254 | 2 | 44.50 | 3.70 | **0.033** |
| | | time point:site | 0.165 | 6 | 76.00 | 0.80 | 0.573 |
| | *Pocillopora* | time point | 1.195 | 3 | 87.24 | 19.44 | **<0.001** |
| | | site | 0.233 | 2 | 34.76 | 5.68 | **0.007** |
| | | time point:site | 0.512 | 6 | 86.86 | 4.17 | **0.001** |
| | *Porites* | time point | 0.635 | 3 | 102.48 | 3.03 | **0.033** |
| | | site | 0.701 | 2 | 41.66 | 41.66 | **0.011** |
| | | time point:site | 1.036 | 6 | 102.27 | 2.47 | **0.028** |
| Respiration (RD) (µmol O2 cm-1 h-1) | *Acropora* | time point | 0.368 | 3 | 69.52 | 13.91 | **<0.001** |
| | | site | 0.134 | 2 | 31.74 | 7.57 | **0.002** |
| | | time point:site | 0.067 | 6 | 67.42 | 1.26 | 0.286 |

| | | | | | | | |
|---|---|---|---|---|---|---|---|
| | *Pocillopora* | time point | 0.015 | 3 | 91.24 | 1.29 | 0.281 |
| | | site | 0.045 | 2 | 39.58 | 5.77 | **0.006** |
| | | time point:site | 0.113 | 6 | 90.90 | 4.86 | **<0.001** |
| | *Porites* | time point | 0.553 | 3 | 128.00 | 10.89 | **<0.001** |
| | | site | 0.114 | 2 | 128.00 | 3.36 | **0.038** |
| | | time point:site | 0.092 | 6 | 128.00 | 0.91 | 0.490 |
| Calcification (µmol CaCO3 mg cm-2 h-1) | *Acropora* | time point | 0.808 | 3 | 78.38 | 36.08 | **<0.001** |
| | | site | 0.034 | 2 | 50.04 | 2.29 | 0.111 |
| | | time point:site | 0.187 | 6 | 77.14 | 4.17 | **0.001** |
| | *Pocillopora* | time point | 0.429 | 3 | 101.71 | 11.46 | **<0.001** |
| | | site | 0.014 | 2 | 45.61 | 0.56 | 0.575 |
| | | time point:site | 0.152 | 6 | 101.33 | 2.03 | 0.068 |
| | *Porites* | time point | 7.236 | 3 | 128.00 | 63.45 | **<0.001** |
| | | site | 0.266 | 2 | 128.00 | 3.51 | **0.033** |
| | | time point:site | 0.655 | 6 | 128.00 | 2.87 | **0.012** |

**Table S8. Univariate linear mixed effect model analysis of symbiont responses.** All responses were log+1 transformed. Linear mixed effect models were run for each genus and included time point, site, and their interactions as main effects with colony nested within holobiont (for *Pocillopora* and *Porites* only) as a random intercept. Significance determined with Type III analysis of variance tests. SS = sum of squares; Num DF = numerator degrees of freedom; Den DF = denominator degrees of freedom. Bold indicates P<0.05. PAR indicates photosynthetically active irradiance. AFDW indicates ash-free dry weight. Holobiont indicates host haplotype and associated symbionts.

| Response | Genus | Main Effect | SS | Num DF | Den DF | F value | P-value |
|---|---|---|---|---|---|---|---|
| Symbiont Cell Density (cells mg AFDW-1) | *Acropora* | time point | 23.200 | 3 | 92.00 | 64.26 | **<0.001** |
| | | site | 0.173 | 2 | 92.00 | 0.72 | 0.489 |
| | | time point:site | 2.154 | 6 | 92.00 | 2.98 | **0.010** |
| | *Pocillopora* | time point | 17.250 | 3 | 86.31 | 47.16 | **<0.001** |
| | | site | 0.549 | 2 | 37.43 | 2.17 | 0.128 |
| | | time point:site | 3.214 | 6 | 86.01 | 4.39 | **0.001** |
| | *Porites* | time point | 10.716 | 3 | 88.10 | 37.05 | **<0.001** |
| | | site | 1.614 | 2 | 26.18 | 8.37 | **0.002** |
| | | time point:site | 1.817 | 6 | 87.97 | 3.14 | **0.008** |
| Symbiont Biomass (mg AFDW cm-2) | *Acropora* | time point | 0.230 | 3 | 83.07 | 4.19 | **0.008** |
| | | site | 0.156 | 2 | 49.10 | 4.25 | **0.020** |
| | | time point:site | 0.247 | 6 | 81.77 | 2.25 | **0.047** |
| | *Pocillopora* | time point | 0.723 | 3 | 91.24 | 17.43 | **<0.001** |
| | | site | 0.104 | 2 | 37.54 | 3.74 | **0.033** |
| | | time point:site | 0.157 | 6 | 90.93 | 1.89 | 0.092 |
| | *Porites* | time point | 0.809 | 3 | 97.52 | 2.42 | 0.071 |
| | | site | 0.596 | 2 | 37.74 | 2.67 | 0.082 |
| | | time point:site | 0.604 | 6 | 97.26 | 0.90 | 0.495 |
| Total Chlorophyll (chl a + chl c2) (µg chl mg AFDW-1) | *Acropora* | time point | 3.481 | 3 | 92.00 | 11.52 | **<0.001** |
| | | site | 0.694 | 2 | 92.00 | 3.44 | **0.036** |
| | | time point:site | 1.926 | 6 | 92.00 | 3.19 | **0.007** |
| | *Pocillopora* | time point | 2.901 | 3 | 122.00 | 18.82 | **<0.001** |
| | | site | 0.878 | 2 | 122.00 | 8.54 | **<0.001** |
| | | time point:site | 1.660 | 6 | 122.00 | 5.39 | **<0.001** |
| | *Porites* | time point | 2.337 | 3 | 128.00 | 10.30 | **<0.001** |
| | | site | 1.699 | 2 | 128.00 | 11.23 | **<0.001** |
| | | time point:site | 1.045 | 6 | 128.00 | 2.30 | **0.038** |
| Total Cell-Specific Chlorophyll (µg cell-1) | *Acropora* | time point | 5.37E-10 | 3 | 7.13 | 13.12 | **0.003** |
| | | site | 6.42E-11 | 2 | 6.18 | 2.35 | 0.174 |
| | | time point:site | 7.13E-11 | 6 | 7.13 | 0.87 | 0.558 |

| | | | | | | | |
|---|---|---|---|---|---|---|---|
| | *Pocillopora* | time point | 1.04E-09 | 3 | 14.22 | 5.52 | **0.010** |
| | | site | 3.78E-10 | 2 | 28.70 | 2.99 | 0.066 |
| | | time point:site | 6.77E-10 | 6 | 14.19 | 1.79 | 0.173 |
| | *Porites* | time point | 4.12E-10 | 3 | 11.01 | 5.28 | **0.017** |
| | | site | 1.23E-10 | 2 | 10.95 | 2.37 | 0.140 |
| | | time point:site | 2.89E-10 | 6 | 11.01 | 1.85 | 0.179 |
| Symbiont : Host Biomass | *Acropora* | time point | 0.074 | 3 | 8.78 | 8.78 | **<0.001** |
| | | site | 0.020 | 2 | 3.62 | 3.62 | **0.034** |
| | | time point:site | 0.039 | 6 | 2.30 | 2.30 | **0.043** |
| | *Pocillopora* | time point | 0.162 | 3 | 121.02 | 30.46 | **<0.001** |
| | | site | 0.007 | 2 | 121.30 | 1.98 | 0.143 |
| | | time point:site | 0.029 | 6 | 121.02 | 2.73 | **0.016** |
| | *Porites* | time point | 0.039 | 3 | 95.67 | 5.57 | **0.001** |
| | | site | 0.003 | 2 | 42.14 | 0.61 | 0.550 |
| | | time point:site | 0.007 | 6 | 95.58 | 0.49 | 0.815 |
| Maximal Photosynthesis (PMAX) (µmol O2 cm-1 h-1) | *Acropora* | time point | 0.468 | 3 | 54.40 | 12.19 | **<0.001** |
| | | site | 0.078 | 2 | 36.82 | 3.03 | 0.060 |
| | | time point:site | 0.172 | 6 | 53.36 | 2.25 | 0.053 |
| | *Pocillopora* | time point | 0.211 | 3 | 95.86 | 7.29 | **<0.001** |
| | | site | 0.059 | 2 | 28.21 | 3.04 | 0.064 |
| | | time point:site | 0.054 | 6 | 95.49 | 0.93 | 0.479 |
| | *Porites* | time point | 0.582 | 3 | 128.00 | 4.93 | **0.003** |
| | | site | 0.309 | 2 | 128.00 | 3.94 | **0.022** |
| | | time point:site | 0.216 | 6 | 128.00 | 0.92 | 0.486 |
| Apparent Quantum Yield (AQY) | *Acropora* | time point | 3.57E-06 | 3 | 44.82 | 2.79 | 0.051 |
| | | site | 1.21E-06 | 2 | 39.70 | 1.42 | 0.255 |
| | | time point:site | 3.14E-06 | 6 | 44.55 | 1.22 | 0.312 |
| | *Pocillopora* | time point | 2.64E-05 | 3 | 92.35 | 11.47 | **<0.001** |
| | | site | 4.66E-06 | 2 | 34.09 | 3.04 | 0.061 |
| | | time point:site | 6.13E-06 | 6 | 91.88 | 1.33 | 0.250 |
| | *Porites* | time point | 1.53E-04 | 3 | 127.07 | 6.36 | **<0.001** |
| | | site | 4.08E-05 | 2 | 103.25 | 2.54 | 0.840 |
| | | time point:site | 6.15E-05 | 6 | 127.04 | 1.28 | 0.272 |
| Saturating Irradiance (IK) (PAR) | *Acropora* | time point | 1.469 | 3 | 59.57 | 7.34 | **<0.001** |
| | | site | 0.117 | 2 | 44.49 | 0.88 | 0.423 |
| | | time point:site | 0.570 | 6 | 58.71 | 1.42 | 0.221 |
| | *Pocillopora* | time point | 7.568 | 3 | 88.58 | 24.84 | **<0.001** |
| | | site | 0.237 | 2 | 28.78 | 1.17 | 0.325 |

| | | | | | | | |
|---|---|---|---|---|---|---|---|
| | | time point:site | 0.543 | 6 | 88.04 | 0.89 | 0.505 |
| | *Porites* | time point | 5.478 | 3 | 127.08 | 16.34 | **<0.001** |
| | | site | 1.717 | 2 | 101.78 | 7.68 | **<0.001** |
| | | time point:site | 2.164 | 6 | 127.04 | 3.23 | **0.006** |
| Compensation Irradiance (IC) (PAR) | *Acropora* | time point | 7.013 | 3 | 70.81 | 12.66 | **<0.001** |
| | | site | 2.440 | 2 | 28.10 | 6.61 | **0.004** |
| | | time point:site | 2.388 | 6 | 68.35 | 2.16 | 0.058 |
| | *Pocillopora* | time point | 2.659 | 3 | 94.09 | 4.91 | **0.003** |
| | | site | 0.194 | 2 | 25.57 | 0.54 | 0.592 |
| | | time point:site | 4.822 | 6 | 93.54 | 4.46 | **<0.001** |
| | *Porites* | time point | 3.378 | 3 | 91.92 | 8.72 | **<0.001** |
| | | site | 0.669 | 2 | 28.52 | 2.59 | 0.093 |
| | | time point:site | 2.930 | 6 | 91.71 | 3.78 | **0.002** |

**Table S9. Permutational multivariate analysis of variance of physiology.** PERMANOVA analyses conducted for each species and biological level (combined responses, host, and symbiont) separately. Time point, site, and their interaction were included as main effects. Holobiont identity (i.e., host haplotype and associated symbiont communities) was also included in *Pocillopora* and *Porites* PERMANOVA models. Omega R2 indicates R2 corrected for degrees of freedom. Bold indicates P<0.05. DF = degrees of freedom; SS = sum of squares. Tests run with 999 permutations.

| Genus | Biological Level | Main Effect | DF | SS | R2 | Omega R2 | F value | P-value |
|---|---|---|---|---|---|---|---|---|
| ***Acropora*** | Combined | time point | 3 | 423.98 | 0.30 | 0.30 | 15.67 | **0.001** |
| | | site | 2 | 95.01 | 0.07 | 0.08 | 5.27 | **0.001** |
| | | time point:site | 6 | 83.2 | 0.06 | 0.03 | 1.54 | **0.025** |
| | | residuals | 90 | 811.91 | 0.57 | | | |
| | Host | time point | 3 | 196.82 | 0.38 | 0.39 | 23.65 | **0.001** |
| | | site | 2 | 35.68 | 0.07 | 0.09 | 6.43 | **0.001** |
| | | time point:site | 6 | 27.3 | 0.05 | 0.04 | 1.64 | **0.033** |
| | | residuals | 92 | 255.21 | 0.50 | | | |
| | Symbiont | time point | 3 | 255.03 | 0.26 | 0.25 | 13.68 | **0.001** |
| | | site | 2 | 60.94 | 0.06 | 0.07 | 4.90 | **0.001** |
| | | time point:site | 6 | 61.66 | 0.06 | 0.03 | 1.65 | **0.021** |
| | | residuals | 100 | 621.37 | 0.62 | | | |
| ***Pocillopora*** | Combined | holobiont | 1 | 67.75 | 0.04 | 0.16 | 7.19 | **0.001** |
| | | time point | 3 | 442.85 | 0.24 | 0.19 | 15.67 | **0.001** |
| | | site | 2 | 69.74 | 0.04 | 0.04 | 3.70 | **0.001** |
| | | time point:site | 6 | 137.41 | 0.07 | 0.04 | 2.43 | **0.001** |
| | | residuals | 120 | 1130.25 | 0.61 | | | |
| | Host | holobiont | 1 | 12.43 | 0.02 | 0.02 | 3.63 | **0.005** |
| | | time point | 3 | 173.34 | 0.24 | 0.25 | 16.88 | **0.001** |
| | | site | 2 | 22.18 | 0.03 | 0.03 | 3.24 | **0.002** |
| | | time point:site | 6 | 61.73 | 0.09 | 0.08 | 3.01 | **0.001** |
| | | residuals | 133 | 455.33 | 0.63 | | | |
| | Symbiont | holobiont | 1 | 45.11 | 0.04 | 0.04 | 7.15 | **0.001** |
| | | time point | 3 | 282.66 | 0.22 | 0.22 | 14.94 | **0.001** |
| | | site | 2 | 39.87 | 0.03 | 0.03 | 3.16 | **0.003** |
| | | time point:site | 6 | 98.76 | 0.08 | 0.06 | 2.61 | **0.001** |
| | | residuals | 133 | 838.62 | 0.64 | | | |

| | | | | | | | | |
|---|---|---|---|---|---|---|---|---|
| ***Porites*** | Combined | holobiont | 1 | 258.36 | 0.13 | 0.04 | 28.05 | **0.001** |
| | | time point | 3 | 329.09 | 0.17 | 0.25 | 11.91 | **0.001** |
| | | site | 2 | 75.05 | 0.04 | 0.04 | 4.08 | **0.001** |
| | | time point:site | 6 | 108.94 | 0.06 | 0.06 | 1.97 | **0.001** |
| | | residuals | 126 | 1160.55 | 0.60 | | | |
| | Host | holobiont | 1 | 92.46 | 0.13 | 0.17 | 29.55 | **0.001** |
| | | time point | 3 | 150.69 | 0.22 | 0.24 | 16.05 | **0.001** |
| | | site | 2 | 22.42 | 0.03 | 0.04 | 3.58 | **0.002** |
| | | time point:site | 6 | 33.90 | 0.05 | 0.03 | 1.81 | **0.012** |
| | | residuals | 128 | 400.53 | 0.57 | | | |
| | Symbiont | holobiont | 1 | 186.77 | 0.13 | 0.16 | 30.78 | **0.001** |
| | | time point | 3 | 202.11 | 0.14 | 0.16 | 11.1 | **0.001** |
| | | site | 2 | 64.47 | 0.05 | 0.05 | 5.31 | **0.001** |
| | | time point:site | 6 | 76.75 | 0.05 | 0.04 | 2.11 | **0.001** |
| | | residuals | 144 | 873.9 | 0.62 | | | |

**Table S10. Permutational multivariate analysis of dispersion.** PERMDISP analyses conducted for each species, biological level (combined responses, host, symbiont), and effects of site, time, and holobiont separately. Bold indicates P<0.05. DF = degrees of freedom; SS = sum of squares. Holobiont refers to host haplotype and associated symbionts.

| Genus | Biological Level | Effect | DF | SS | F value | P-value |
|---|---|---|---|---|---|---|
| ***Acropora*** | Combined | site | 2 | 1.68 | 5.99 | **0.004** |
| | | time point | 3 | 0.83 | 2.69 | 0.051 |
| | Host | site | 2 | 0.16 | 4.41 | **0.015** |
| | | time point | 3 | 0.02 | 0.34 | 0.794 |
| | Symbiont | site | 2 | 2.07 | 6.56 | **0.002** |
| | | time point | 3 | 1.34 | 3.76 | **0.013** |
| ***Pocillopora*** | Combined | site | 2 | 0.53 | 2.46 | 0.089 |
| | | holobiont | 1 | 0.06 | 0.52 | 0.471 |
| | | time point | 3 | 2.09 | 7.47 | **<0.001** |
| | Host | site | 2 | 0.08 | 1.70 | 0.186 |
| | | holobiont | 1 | 0.02 | 0.75 | 0.389 |
| | | time point | 3 | 0.15 | 2.81 | **0.042** |
| | Symbiont | site | 2 | 0.77 | 3.49 | **0.033** |
| | | holobiont | 1 | 0.06 | 0.50 | 0.481 |
| | | time point | 3 | 1.90 | 6.41 | **<0.001** |
| ***Porites*** | Combined | site | 2 | 2.35 | 8.87 | **<0.001** |
| | | holobiont | 1 | 0.34 | 2.53 | 0.114 |
| | | time point | 3 | 1.01 | 2.34 | 0.076 |
| | Host | site | 2 | 0.53 | 2.46 | 0.089 |
| | | holobiont | 1 | 0.15 | 4.53 | **0.035** |
| | | time point | 3 | 0.027 | 2.68 | 0.050 |
| | Symbiont | site | 2 | 2.43 | 8.73 | **<0.001** |
| | | holobiont | 1 | 0.84 | 2.15 | 0.097 |
| | | time point | 3 | 0.25 | 1.86 | 0.175 |

**Table S11. Significance of variance explained by main effects on multivariate physiology.** Variance partitoning analyses conducted for each genus, biological level (combined responses, host, symbiont), and effects of site, time, and holobiont identity (i.e. coral host haplotype and associated symbiont community). Significance of individual main effects tested by testing each main effect while controlling for the other main effects in turn. F test statistics and P-values for each main effect determined from ANOVA-like permutation analyses of partial redundancy analyses (RDA). DF = degrees of freedom. Holobiont main effect refers to the host genetic haplotype and associated symbiont communities. Combined responses are all host and symbiont responses analyzed together. Bold indicates P<0.05.

| Genus | Biological Level | Effect Tested | Effect | DF | Variance | F | P-value |
|---|---|---|---|---|---|---|---|
| ***Acropora*** | Combined | time | site | 3 | 0.40 | 22.23 | **0.001** |
| | | site | time | 2 | 0.04 | 3.58 | **0.001** |
| | Host | time | site | 3 | 0.02 | 14.50 | **0.001** |
| | | site | time | 2 | 0.01 | 7.01 | **0.001** |
| | Symbiont | time | site | 3 | 0.39 | 25.01 | **0.001** |
| | | site | time | 2 | 0.03 | 3.24 | **0.004** |
| ***Pocillopora*** | Combined | time | site | 3 | 0.27 | 18.39 | **0.001** |
| | | time | holobiont | 3 | 0.27 | 18.12 | **0.001** |
| | | site | time | 2 | 0.02 | 2.05 | 0.050 |
| | | site | holobiont | 2 | 0.02 | 1.69 | 0.131 |
| | | holobiont | time | 1 | 0.01 | 1.39 | 0.214 |
| | | holobiont | site | 1 | 0.01 | 1.53 | 0.186 |
| | Host | time | site | 3 | 0.01 | 12.28 | **0.001** |
| | | time | holobiont | 3 | 0.01 | 11.92 | **0.001** |
| | | site | time | 2 | 0.003 | 4.97 | **0.002** |
| | | site | holobiont | 2 | 0.003 | 3.32 | **0.015** |
| | | holobiont | time | 1 | 0.001 | 2.6 | 0.069 |
| | | holobiont | site | 1 | 0.00 | 0.41 | 0.711 |
| | Symbiont | time | site | 3 | 0.24 | 19.35 | **0.001** |
| | | time | holobiont | 3 | 0.24 | 19.40 | **0.001** |
| | | site | time | 2 | 0.01 | 1.54 | 0.141 |
| | | site | holobiont | 2 | 0.01 | 1.25 | 0.273 |
| | | holobiont | time | 1 | 0.01 | 1.68 | 0.150 |
| | | holobiont | site | 1 | 0.01 | 1.26 | 0.290 |

| | | | | | | | |
|---|---|---|---|---|---|---|---|
| ***Porites*** | Combined | time | site | 3 | 0.24 | 13.68 | **0.001** |
| | | time | holobiont | 3 | 0.24 | 14.43 | **0.001** |
| | | site | time | 2 | 0.13 | 11.53 | **0.001** |
| | | site | holobiont | 2 | 0.06 | 4.64 | **0.001** |
| | | holobiont | time | 1 | 0.14 | 25.94 | **0.001** |
| | | holobiont | site | 1 | 0.07 | 10.18 | **0.001** |
| | Host | time | site | 3 | 0.06 | 19.65 | **0.001** |
| | | time | holobiont | 3 | 0.06 | 19.20 | **0.001** |
| | | site | time | 2 | 0.01 | 5.79 | **0.001** |
| | | site | holobiont | 2 | 0.00 | 1.71 | 0.125 |
| | | holobiont | time | 1 | 0.01 | 8.41 | **0.002** |
| | | holobiont | site | 1 | 0.00 | 1.61 | 0.187 |
| | Symbiont | time | site | 3 | 0.17 | 15.06 | **0.001** |
| | | time | holobiont | 3 | 0.17 | 15.45 | **0.001** |
| | | site | time | 2 | 0.12 | 15.23 | **0.001** |
| | | site | holobiont | 2 | 0.05 | 5.98 | **0.001** |
| | | holobiont | time | 1 | 0.13 | 33.23 | **0.001** |
| | | holobiont | site | 1 | 0.06 | 13.57 | **0.001** |

**Table S12. Evaluation of random effects in univariate linear mixed effect model analysis of host responses.** ANOVA-like analysis of random effects of linear mixed effect models on each response using single term deletions. LogLik = log likelihood; AIC = Akaike Information Criterion; LRT = likelihood ratio test; DF = degrees of freedom. All responses were log+1 transformed in linear mixed effect model analysis. Linear mixed effect models were conducted for each genus and included time point, site, and their interactions as main effects with colony nested within holobiont (for *Porites* and *Pocillopora* only) as random effects. Bold indicates P<0.05. CRE indicates copper reducing elements. AFDW indicates ash-free dry weight. Holobiont indicates host haplotype and associated symbionts.

| Response | Genus | Random Effect | LogLik | AIC | LRT | DF | P-value |
|---|---|---|---|---|---|---|---|
| Antioxidant Capacity (µmol CRE mg AFDW-1) | *Acropora* | colony | 171.27 | -316.53 | 3.60 | 1 | 0.058 |
| | *Pocillopora* | colony:holobiont | 288.63 | -549.27 | 0.00 | 1 | 1.000 |
| | | holobiont | 288.63 | -549.27 | 0.00 | 1 | 1.000 |
| | *Porites* | colony:holobiont | 172.34 | -316.69 | 21.25 | 1 | **<0.001** |
| | | holobiont | 176.27 | -324.54 | 13.40 | 1 | **<0.001** |
| Host Soluble Protein (mg protein mg AFDW-1) | *Acropora* | colony | 120.38 | -214.76 | 0.50 | 1 | 0.480 |
| | *Pocillopora* | colony:holobiont | 220.47 | -412.95 | 0.00 | 1 | 1.000 |
| | | holobiont | 219.68 | -411.35 | 1.59 | 1 | 0.207 |
| | *Porites* | colony:holobiont | 149.54 | -271.09 | 6.66 | 1 | **0.010** |
| | | holobiont | 146.16 | -264.33 | 13.42 | 1 | **<0.001** |
| Host Biomass (mg AFDW cm-2) | *Acropora* | colony | 9.18 | 7.63 | 0.33 | 1 | 0.568 |
| | *Pocillopora* | colony:holobiont | 37.03 | -46.06 | 3.32 | 1 | 0.069 |
| | | holobiont | 38.69 | -49.38 | 0.00 | 1 | 1.000 |
| | *Porites* | colony:holobiont | -28.16 | 81.32 | 0.19 | 1 | 0.666 |
| | | holobiont | -28.07 | 84.14 | 0.00 | 1 | 1.000 |
| Respiration (RD) (µmol O2 cm-1 h-1) | *Acropora* | colony | 72.00 | -118.00 | 0.16 | 1 | 0.690 |
| | *Pocillopora* | colony:holobiont | 136.76 | -245.53 | 5.76 | 1 | **0.016** |
| | | holobiont | 139.64 | -251.28 | 0.00 | 1 | 1.000 |
| | *Porites* | colony:holobiont | 64.83 | -101.66 | 0.00 | 1 | 1.000 |
| | | holobiont | 64.83 | -101.66 | 0.00 | 1 | 1.000 |
| Calcification (µmol CaCO3 mg cm-2 h-1) | *Acropora* | colony | 77.98 | -129.08 | 0.88 | 1 | 0.348 |
| | *Pocillopora* | colony:holobiont | 78.08 | -128.15 | 0.27 | 1 | 0.602 |
| | | holobiont | 78.21 | -128.42 | 0.00 | 1 | 1.000 |
| | *Porites* | colony:holobiont | 12.98 | 2.04 | 0.00 | 1 | 1.000 |
| | | holobiont | 12.98 | 2.04 | 0.00 | 1 | 1.000 |

**Table S13. Evaluation of random effects in univariate linear mixed effect model analysis of symbiont responses.** ANOVA-like analysis of random effects of linear mixed effect models on each response using single term deletions. LogLik = log likelihood; AIC = Akaike Information Criterion; LRT = likelihood ratio test; DF = degrees of freedom. All responses were log+1 transformed in linear mixed effect model analysis. Linear mixed effect models were conducted for each genus and included time point, site, and their interactions as main effects with colony nested within holobiont (for *Porites* and *Pocillopora* only) as random effects. Bold indicates P<0.05. PAR indicates photosynthetically active irradiance. AFDW indicates ash-free dry weight. Holobiont indicates host haplotype and associated symbionts.

| Response | Genus | Random Effect | LogLik | AIC | LRT | DF | P-value |
|---|---|---|---|---|---|---|---|
| Symbiont Cell Density (cells mg AFDW-1) | *Acropora* | colony | -45.41 | 116.81 | 7.11E-14 | 1 | 1.000 |
| | *Pocillopora* | colony:holobiont | -81.54 | 191.07 | 9.71 | 1 | **0.002** |
| | | holobiont | -76.68 | 181.36 | 0.00 | 1 | 1.000 |
| | *Porites* | colony:holobiont | -50.42 | 128.84 | 0.09 | 1 | 0.760 |
| | | holobiont | -59.09 | 146.18 | 17.43 | 1 | **<0.001** |
| Symbiont Biomass (mg AFDW cm-2) | *Acropora* | colony | 40.98 | -55.96 | 0.00 | 1 | 0.952 |
| | *Pocillopora* | colony:holobiont | 60.37 | -92.75 | 3.48 | 1 | 0.062 |
| | | holobiont | 52.79 | -77.57 | 18.65 | 1 | **<0.001** |
| | *Porites* | colony:holobiont | -63.92 | 155.83 | 1.20 | 1 | 0.273 |
| | | holobiont | -69.74 | 167.47 | 12.84 | 1 | **<0.001** |
| Total Chlorophyll (chl a + chl c2) (µg chl mg AFDW-1) | *Acropora* | colony | -37.21 | 100.42 | 0.00 | 1 | 1.000 |
| | *Pocillopora* | colony:holobiont | -6.41 | 40.81 | 0.00 | 1 | 1.000 |
| | | holobiont | -6.41 | 40.81 | 0.00 | 1 | 1.000 |
| | *Porites* | colony:holobiont | -31.07 | 90.15 | 0.00 | 1 | 1.000 |
| | | holobiont | -31.07 | 90.15 | -6.39E-14 | 1 | 1.000 |
| Total Cell-Specific Chlorophyll (chl a + chl c2) (µg cell-1) | *Acropora* | colony | 1001.20 | -1976.4 | 1.03 | 1 | 0.311 |
| | *Pocillopora* | colony:holobiont | 1237.20 | -2446.4 | 7.26 | 1 | **0.007** |
| | | holobiont | 1240.80 | -2453.7 | 0.00 | 1 | 1.000 |
| | *Porites* | colony:holobiont | 1362.40 | -2696.8 | 0.00 | 1 | 1.000 |
| | | holobiont | 1359.80 | -2691.7 | 5.11 | 1 | **0.024** |
| Symbiont : Host Biomass | *Acropora* | colony | 123.24 | -220.49 | 0.67 | 1 | 0.415 |
| | *Pocillopora* | colony:holobiont | 197.07 | -366.14 | 0.00 | 1 | 1.000 |
| | | holobiont | 183.97 | -339.93 | 26.21 | 1 | **<0.001** |
| | *Porites* | colony:holobiont | 167.58 | -307.16 | 10.93 | 1 | **<0.001** |
| | | holobiont | 166.76 | -305.53 | 12.56 | 1 | **<0.001** |

| | | | | | | | |
|---|---|---|---|---|---|---|---|
| Maximal Photosynthesis (PMAX) (µmol O2 cm-1 h-1) | *Acropora* | colony | 39.79 | -53.75 | 4.84 | 1 | **0.028** |
| | *Pocillopora* | colony:holobiont | 94.64 | -161.27 | 0.10 | 1 | 0.757 |
| | | holobiont | 94.68 | -161.36 | 0.01 | 1 | 0.931 |
| | *Porites* | colony:holobiont | 10.87 | 6.27 | 0.00 | 1 | 1.000 |
| | | holobiont | 10.87 | 6.27 | 0.00 | 1 | 1.000 |
| Apparent Quantum Yield (AQY) | *Acropora* | colony | 479.22 | -932.44 | 17.97 | 1 | **<0.001** |
| | *Pocillopora* | colony:holobiont | 673.54 | -1319.1 | 0.32 | 1 | 0.571 |
| | | holobiont | 673.70 | -1319.4 | 0.00 | 1 | 1.000 |
| | *Porites* | colony:holobiont | 553.83 | -1079.7 | 0.00 | 1 | 1.000 |
| | | holobiont | 552.73 | -1077.5 | 2.21 | 1 | 0.137 |
| Saturating Irradiance (IK) (PAR) | *Acropora* | colony | -39.41 | 104.82 | 6.87 | 1 | **0.009** |
| | *Pocillopora* | colony:holobiont | -49.66 | 127.31 | 0.04 | 1 | 0.840 |
| | | holobiont | -49.63 | 127.27 | 0.00 | 1 | 1.000 |
| | *Porites* | colony:holobiont | -56.80 | 141.6 | 0.00 | 1 | 1.000 |
| | | holobiont | -57.85 | 143.7 | 2.10 | 1 | 0.147 |
| Compensation Irradiance (IC) (PAR) | *Acropora* | colony | -65.84 | 157.69 | 0.01 | 1 | 0.918 |
| | *Pocillopora* | colony:holobiont | -84.79 | 197.59 | 0.04 | 1 | 0.843 |
| | | holobiont | -84.77 | 197.55 | 0.00 | 1 | 0.967 |
| | *Porites* | colony:holobiont | -67.02 | 162.04 | 0.00 | 1 | 0.954 |
| | | holobiont | -71.99 | 171.97 | 9.93 | 1 | **0.002** |

**Table S14. Redundancy analysis of variance constrained in symbiont and host physiology by environmental characteristics.** P-values of each term from redundancy analysis (RDA) and model analysis of variance constrained in host and symbiont of each genus by host and symbiont responses with holobiont identity included in models for each biological level. Main effects were mean light (solar radiance in kWh m-2) and mean temperature (°C) that were scaled for analysis. Gray boxes indicate no significant effect. Bold indicates P<0.05.

| Biological Level | Genus | Main Effect | DF | Variance | F | P-value |
|---|---|---|---|---|---|---|
| Host Responses | *Acropora* | light | 1 | 0.66 | 13.04 | **0.001** |
| | | temperature | 1 | 0.27 | 5.41 | **0.002** |
| | *Pocillopora* | light | 1 | 0.01 | 9.73 | **0.001** |
| | | temperature | 1 | 0.01 | 19.00 | **0.001** |
| | *Porites* | light | 1 | 0.01 | 4.18 | **0.007** |
| | | temperature | 1 | 0.05 | 30.58 | **0.001** |
| Symbiont Responses | *Acropora* | light | 1 | 0.30 | 52.63 | **0.001** |
| | | temperature | 1 | 0.11 | 18.75 | **0.001** |
| | *Pocillopora* | light | 1 | 0.19 | 38.58 | **0.001** |
| | | temperature | 1 | 0.06 | 12.04 | **0.001** |
| | *Porites* | light | 1 | 0.11 | 20.67 | **0.001** |
| | | temperature | 1 | 0.05 | 9.17 | **0.001** |

**Table S15. Redundancy analysis of variance constrained in symbiont ITS2 communities by host and symbiont responses.** Distance-based RDA analysis of variance constrained in symbiont communities of each species by host and symbiont responses. Variance constrained by host and symbiont responses on symbiont communities in each species expresed as a percentage of variance in symbiont ITS2 community explained by significant models. Bold indicates P<0.05. DF = degrees of freedom/number of responses tested at each biological level; SS = sum of squares.

| Genus | (Biological | DF | SS | F value | P-value | Community | Community by Model |
|---|---|---|---|---|---|---|---|
| ***Acropora*** | Host | 5 | 0.38 | 0.64 | 0.751 | | |
| | Symbiont | 9 | 0.90 | 0.84 | 0.624 | | |
| ***Pocillopora*** | Host | 5 | 1.29 | 0.65 | 0.886 | 2.64 | 0 |
| | Symbiont | 9 | 6.88 | 2.11 | **0.001** | 14.09 | 7.43 |
| ***Porites*** | Host | 5 | 6.68 | 4.26 | **0.001** | 14.17 | 10.84 |
| | Symbiont | 9 | 10.66 | 4.05 | **0.001** | 22.58 | 17.01 |

**Table S16. Redundancy analysis of variance constrained in symbiont ITS2 communities by host and symbiont responses at the genus level.** P-values of each term from distance-based RDA (dbRDA) analysis of variance constrained in symbiont communities of each genus by host and symbiont responses with holobiont identity included in models for each biological level. Significance of variance explained in symbiont communities by each host and symbiont response as analyzed with dbRDA models. Gray boxes indicate no significant effect. Bold indicates $P<0.05$.

| | ***Acropora*** | ***Pocillopora*** | ***Porites*** |
|---|---|---|---|
| **Host Responses** | | | |
| Antioxidant capacity | 0.277 | 0.992 | **0.001** |
| Host biomass | 0.366 | 0.791 | **0.002** |
| Host protein | 0.878 | 0.826 | 0.077 |
| Respiration | 0.543 | 0.096 | 0.428 |
| Calcification | 0.671 | 0.826 | 0.269 |
| **Symbiont Responses** | | | |
| Cell density | 0.145 | 0.514 | **0.002** |
| Symbiont biomass | 0.512 | **0.014** | **0.001** |
| Total chlorophyll | 0.684 | **0.001** | 0.423 |
| Total chlorophyll per cell | 0.402 | **0.046** | 0.101 |
| PMAX | 0.832 | 0.921 | **0.038** |
| AQY | 0.645 | 0.757 | **0.002** |
| IK | 0.235 | 0.254 | 0.354 |
| IC | 0.159 | **0.024** | 0.214 |
| S:H biomass | 0.767 | **0.033** | 0.438 |
